# *Salmonella* SiiE-mediated apical invasion into colonocytes depends on MUC1 α2,3-linked sialic acids

**DOI:** 10.64898/2026.07.21.739827

**Authors:** Koen C.A.P. Giesbers, Agnes L. Hipgrave Ederveen, Helena Coelho, Liane Z.X. Huang, Albert van Dijk, Bart Westendorp, Jinyi Su, Annemarie Kuipers, Angelina S. Palma, Jos P.M. van Putten, Noortje de Haan, Joanne Donkers, Karin Strijbis

## Abstract

MUC1 is a highly O-glycosylated cell-bound mucin that plays key roles in intestinal mucosal maintenance and microbe–host interactions. The enteropathogen *Salmonella enterica* expresses a giant adhesin SiiE, which mediates interaction with MUC1 and apical invasion of epithelial cells in a sialic acid–dependent manner. Here, we investigated the glycan specificity of the SiiE–MUC1 interaction and the expression of glycosylated MUC1 receptor in advanced intestinal epithelial models. Expression of the SiiE adhesin by *Salmonella* was highest in late logarithmic growth, could be induced by aerobic shock, and was detectable on the bacterial surface and in culture supernatant. Purified SiiE bound multiple *O*-glycan structures in a MUC1 glycopeptide array, including those bearing terminal sialic acids. Single-cell RNA sequencing of human intestinal epithelium showed that high MUC1 expression correlated with expression of ST3GAL and ST6GALNAC sialyltransferases, indicating the potential presence of both α2,3- and α2,6-linked sialylation *in vivo*. In HT29-MTX intestinal cultures, both α2,3- and α2,6-linked sialic acids could be detected on the apical surface and α2,3-sialic acid staining colocalized with MUC1. Mass spectrometry–based O-glycomics demonstrated that MUC1 carried predominantly core 1 and core 2 O-glycans decorated with α2,3-linked sialylation. Removal or blocking of α2,3-linked sialic acids abolished *Salmonella* invasion through the SiiE-MUC1 route. In advanced *ex vivo* cultures of human ileum and colon, MUC1 was detected in the colon, where regions showed positive staining for α2,3-linked sialic acids, but not in the ileum. After infection of the *ex vivo* tissues, *Salmonella* was found in close proximity to α2,3-sialylated colonic MUC1. Together, these findings demonstrate that *Salmonella* SiiE-mediated apical invasion of colonocytes depends on α2,3-sialylated O-glycans on MUC1. In humans, this pathway might be most relevant during *Salmonella* invasion in the colon.

## Introduction

In the gastrointestinal tract, a family of large glycoproteins called mucins play a key role in maintaining intestinal homeostasis and protecting the epithelium from microbial invasion [1]. Secreted gel-forming mucins such as MUC2 are secreted by goblet cells and transmembrane mucins such as MUC1, MUC3, MUC13, and MUC17 are expressed on the apical surface of the different types of intestinal epithelial cells [2]. Expression patterns of transmembrane (TM) mucins are cell- and region-specific and presumably tailored to the specific requirements of each region. TM mucins form a glycocalyx that allows interaction with commensal bacteria, they are important players in epithelial barrier formation against invasive pathogens as they regulate mucosal maintenance [3–5]. Transmembrane mucins are a diverse family with different domain structures and varying lengths. In general, they consist of a highly O*-*glycosylated extracellular domain, a Sperm protein, Enterokinase, and Agrin (SEA) domain, a transmembrane region and a cytoplasmic tail containing potential phosphorylation sites that link to signaling pathways [5]. MUC1 is one of the most well-studied TM mucins and was demonstrated to play a protective role during *Helicobacter pylori* and *Campylobacter jejuni* infection by reducing bacterial attachment, acting as a decoy receptor and modulating NFkB responses of the epithelium [6–8]. However, pathogens can also utilize TM mucins for adhesion or invasion. We demonstrated that *Salmonella* expresses an adhesin that mediates interaction with MUC1 and subsequent apical invasion into intestinal epithelial cells [9].

The extracellular domains (EDs) of mucins are heavily decorated with O-glycan structures, that account for the majority of their molecular mass [5]. The glycosylated ED of MUC1 forms an extended structure that protrudes 200–500 nm from the plasma membrane [10] and mediate interactions with microbes. The MUC1 ED contains a variable number of tandem repeats (VNTR) that are rich in serine and threonine residues on which mucin type O-glycosylation is initiated through the addition of an *N*-acetylgalactosamine (*O*-GalNAc). Glycosyltransferases with different specificities elongate this initial monosaccharide into eight distinct core types. Among these, cores 1 and 2 are the most commonly found on mucins [11]. Human mucin O-glycans are typically capped by fucose or sialic acid, specifically *N*-acetylneuraminic acid (Neu5Ac). Together, these steps can lead to a high diversity of complex O-glycan structures that can be tailored for different functions. The spatial arrangement of these glycan structures forms a so-called ‘’glycan barcode’’ which is an important determinant in microbe-host interactions [12–14].

Sialic acids play an important role in preserving the viscoelastic properties of the mucus layer [15] and in protecting mucins against enzymatic degradation [16]. Due to their surface-exposed position, sialic acids are often the target for microbial adhesins or enzymes. In mucin O*-*glycans, sialic acids are linked to underlying residues via α2,3- and α2,6-linkages. These structural differences are major determinants of the functions of sialic acid structures and their interactions with bacteria and viruses. For instance, α2,6-linked sialylation on MUC2 inhibits proteolytic cleavage by the StcE mucinase expressed by enteropathogenic *E. coli* [17, 18]. Bacterial binding and invasion can also depend on the sialic acid linkage type. *Helicobacter pylori* binds preferentially to α2,3-linked sialic acids with its adhesin SabA in gastric tissue, whereas *Moraxella catarrhalis*, a pathogen involved in pneumonia and COPD exacerbations, binds α2,6-linked sialic acids, likely via the OmpCD protein [19–22]. Human and swine influenza viruses preferentially bind α2,6-linked sialic acids, while avian and equine strains target the α2,3-linked forms [23–25]. Sialic acids also modulate immune activation or suppression depending on their specific modifications and linkages. For example, the macrophage receptor Siglec-1 binds α2,3-linked sialic acids on bacteria to promote phagocytosis, while Siglec-2 on B-cells recognizes α2,6-linked sialic acids (mostly in *cis*) and acts as an inhibitory co-receptor by dampening B-cell receptor signaling upon antigen stimulation [26, 27].

*Salmonella* bacteria are among the leading causes of foodborne disease, responsible for numerous infections and deaths each year [28]. The *Salmonella* genome contains several pathogenicity islands encoding virulence factors that mediate epithelial invasion [29]. *Salmonella* employs its type III secretion system-1 (T3SS-1) to inject effector proteins into host cells, triggering actin remodeling and bacterial uptake [30]. It also employs adhesins such as Rck to invade host cells through a zipper-like mechanism [31]. Another important adhesin is the *Salmonella* giant adhesin SiiE, a 595 kDa protein secreted or retained via the type I secretion system (T1SS) [32]. SiiE contains an impressive amount of 53 lectin-like repeat domains that can interact with different glycan structures including *N*-acetylglucosamine (GlcNAc) and sialic acids (Fig. 1A) [33]. We previously demonstrated that *Salmonella* SiiE can interact with MUC1 in a sialic acid-dependent manner and that this interaction allows the bacteria to invade at the apical surface of intestinal epithelial cells [9]. The occurrence of α2,3-linked and α2,6-linked sialic acids in different regions in the intestine, on specific cell types or on specific mucin proteins is poorly understood. In the current study, we aimed to determine glycan specificity of *Salmonella* SiiE-mediated invasion through MUC1 at the intestinal mucosal surface.

**Figure 1.**
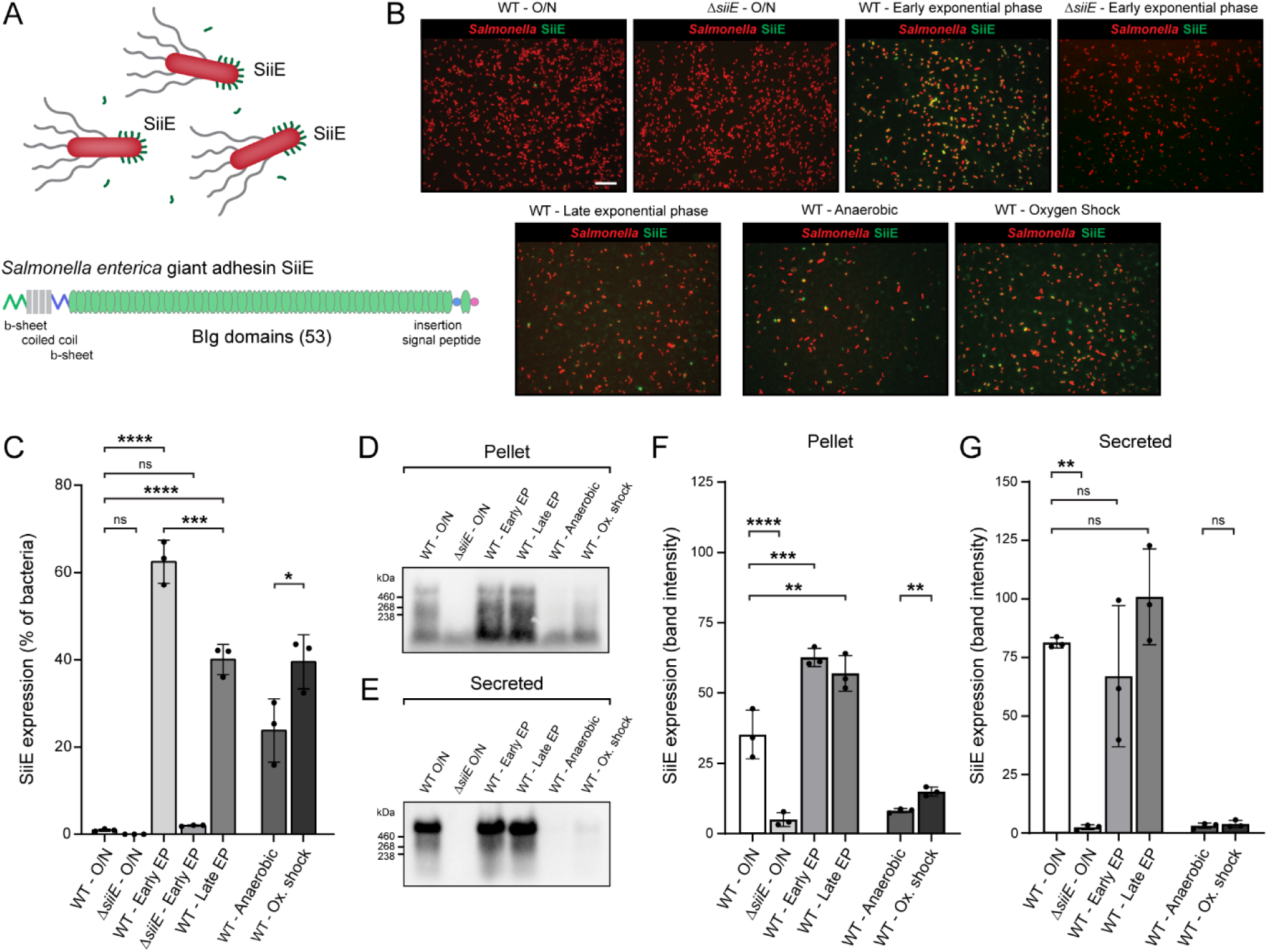
Expression and secretion of *Salmonella* SiiE giant adhesin is dependent of growth phase. A) Schematic representation of the *Salmonella enterica* giant adhesin SiiE expressed on the bacteria. The SiiE adhesin is a 600 kDa multidomain protein with a β-sheet, coiled coil, β-sheet, 53 lectin like Blg domains, insertion domain and signal peptide. Image adapted from Hensel *et al*. 2015 [33]. B) Immunofluorescence of mCherry-positive *Salmonella enterica* serovar Enteritidis (SE) (red) stained for SiiE (green) grown overnight (O/N), to early exponential phase (Early EP), late exponential phase (Late EP), anaerobically or under oxygen shock conditions. Scale bar: 10 µm. C) Quantification of SiiE expression in conditions depicted in B. D, E) Western blot analysis of SiiE expression in bacterial pellet (pellet) and culture supernatant (secreted) during different SE growth conditions. F, G) Quantification of SiiE expression in pellet and supernatant fractions as depicted in D and E. Graphs represent mean ± SD from three independent biological replicates. Statistical test: two-way ANOVA with Tukey’s correction. ns = not significant; * p<0.05; ** p<0.1; *** p<0.001; **** p<0.0001.

## Materials and methods

### Bacterial growth conditions, protein isolation and lysate preparation

*Salmonella enterica* serovar Enteritidis (*S*. Enteritidis) strain 90-13-706 (CVI, Lelystad) and the ΔSiiE deletion mutant were previously described [9]. On day 1, *Salmonella* wild type (WT) and ΔSiiE were streaked on an LB agar plate and incubated overnight at 37°C. The next day, one colony was incubated in 25 mL in a 250 mL Erlenmeyer flask and incubated shaking overnight at 37°C. The next day, bacteria were cultured under diverse growth conditions that were previously established [34]. For overnight (O/N) samples, bacteria were directly harvested. For OD_600_=1 and OD_600_=2 conditions, bacteria were inoculated 1:1000 from overnight cultures in 25 mL LB media in a 250 mL Erlenmeyer flask and incubated shaking at 180 rpm at 37°C until the cultures reached OD_600_ 1 and 2 after 4 to 5.5 hours, respectively. For anaerobic and oxygen shock conditions, bacteria were inoculated 1:1000 in 50 mL in a tightly closed 50 mL Falcon tube, incubated at 37°C without shaking until OD_600_=0.3 was reached after approximately 4 hours. For oxygen shock, the anaerobic culture was followed by transfer to a 250 mL Erlenmeyer flask and shaking at 180 rpm for 15 minutes. All conditions were harvested by centrifugation at 4000 × g for 30 minutes at 4°C. The supernatant fractions were collected by transfer to a fresh tube, and the bacterial pellet stored for further analysis. Bacterial cell pellets were lysed by adding 50-200μL of 2X Laemmli buffer without bromophenol blue (2.5% SDS; 20% glycerol; 120 mM Tris-HCl, pH 6.8) and boiling for 10 min with vortexing every minute. Protein concentrations were measured using the Lowry protein assay [35]. Afterwards, 0.02% of bromophenol blue was added and 30 μg of each sample was loaded on an agarose gel (see below). The culture supernatant fractions were filtered through a 0.22 μm syringe filter and concentrated using a 20 mL 100 kDa centrifugal filter unit (Corning) to about 500 μL for OD_600_=2.0 and 50 μL for OD_600_=0.3 (anaerobic growth condition). Concentrated samples were diluted based on the final volume of supernatant*final OD_600_= factor of dilution. After normalizing the supernatant samples, 2,5 μL of 5X laemmli (5% SDS; 50% glycerol; 250 mM Tris-HCL, pH 6.8; 0.1% bromophenol blue; 5% β-mercaptoethanol) was added to 20 μL of the concentrated supernatant and boiled for 15 minutes at 95°C. 12.5 μL of each sample was loaded on gel.

### Fluorescence in situ hybridization (FISH) of *Salmonella* on Poly-L-lysine slides

*Salmonella* cultures were prepared as described above and diluted to OD_600_=0.1 in Dulbecco’s phosphate-buffered saline (DPBS; D8537, Sigma-Aldrich) and 200 µL aliquots were applied to Poly-L-lysine–coated glass slides (P0425, Sigma-Aldrich) and incubated for 30 min at room temperature (RT) to allow bacterial adherence. Bacteria suspensions were removed, and slides were fixed with 4% paraformaldehyde (PFA) in PBS (J19943, VWR) for 30 min at RT. Fixation was quenched by incubation with 500 µL of 50 mM NH₄Cl in PBS for 10 min at RT, followed by two washes in PBS. Slides were inverted onto 30 µL droplets of hybridization buffer (0.9 M NaCl, 20 mM Tris-HCl pH 7.5, 0.1% SDS (15553027, Invitrogen), and 20% formamide (24311291, VWR) containing 1000 nM of either the EUB338-AF488 probe (5′-GCTGCCTCCCGTAGGAGT-3′, Alexa Fluor 488, Thermo Fisher Scientific), the *Salmonella*-specific Sal3-AF647 probe [36] (5′-AATCACTTCACCTACGTG-3′, Alexa Fluor 647, Thermo Fisher Scientific), or the non-EUB338 control probe (5′-ACTCCTACGGGAGGCAGC-3′, Alexa Fluor 647, Thermo Fisher Scientific). Hybridization was carried out for 2 h and 45 min at 50 °C in a humidified chamber. Following hybridization, slides were immersed in 1 mL hybridization buffer for 15 min at RT, then washed twice with 1 mL PBS supplemented with 0.2% bovine serum albumin (BSA; A7030, Sigma-Aldrich) for 5 min. Slides were rinsed in Milli-Q water, dried, and mounted in ProLong Diamond Antifade Mountant (P36965, Thermo Fisher Scientific). Images were acquired using an EVOS M5000 Imaging System (Thermo Fisher Scientific).

### Protein agarose gels

To detect the high molecular weight SiiE protein, a modified agarose gel electrophoresis protocol, adapted from [37], was employed. A 1.0% agarose gel was prepared by dissolving UltraPure^TM^ agarose (Invitrogen) in 300 mL 1X TAE buffer, boiling the solution, and adding 1% SDS after cooling. Bacterial pellet and normalized supernatant samples were loaded onto the gel, along with a HiMark protein ladder (Thermo Fisher Scientific) and run for 2 hours at 100 V. Proteins were transferred to a PVDF membrane using a Southern blot setup with transfer buffer (0.6 M NaCl, 60 mM sodium citrate; 1 L) overnight. The following day, membranes were blocked with 5% BSA in TSMT buffer (20 mM Tris, 150 mM NaCl, Merck; 1 mM CaCl₂, Sigma-Aldrich; 2 mM MgCl₂, Merck; adjusted to pH 7 with HCl, and 0.1% Tween 20, Sigma-Aldrich) for 1 hour at room temperature (RT). Membranes were then washed with TSMT and incubated with α-SiiE polyclonal rabbit serum (1:2000, a kind gift of Prof. Dr. Michael Hensel, Universität Osnabrück, Germany) in TSMT containing 1% BSA for 1 hour at RT. After another wash with TSMT, membranes were incubated with an α-rabbit IgG secondary antibody (A4914, Sigma-Aldrich) diluted 1:10,000 in TSMT with 1% BSA for 1 hour at RT, followed by 3 washes with TSMT and 1 wash with TSM. Blots were developed using the Clarity Western ECL substrate kit (Bio-Rad) and imaged with a Gel-Doc system (Bio-Rad). Quantification was performed using ImageJ software.

### Bioinformatics single-cell studies

Single-cell gene expression from intestinal epithelial cells was analyzed using a public single-cell RNA-sequencing dataset [38]. The H5AD file containing data from all epithelial cells was downloaded from https://www.gutcellatlas.org and further analyzed in Rstudio using the packages Seurat [39] (https://satijalab.org/seurat/), SeuratData [40] (https://github.com/satijalab/seurat-data) and SeuratDisk (https://github.com/mojaveazure/seurat-disk). Cells from healthy adult subjects were selected and low-quality cells (less than 500 genes or >50% of counts mapping to mitochondrial genes) were removed. Data from the remaining 37,325 cells were then normalized using the SCTransform algorithm [41] (https://github.com/satijalab/sctransform), and dot plots showing the normalized expression by cell type or by intestinal cell type were made. Rare cell types (less than 100 in the dataset) are not shown in the plots. Stacked bar charts were made as following: we assigned cells as positive when at least 1 unique transcript count of the indicated genes was found in the raw count tables.

### Bioinformatics bulk RNAseq HT29-MTX cells

Sialyltransferases gene expression in HT29-MTX was extracted from public dataset (GSE304522) of bulk RNAseq from untreated full confluent HT29-MTX cells cultured for 7 days. Selected sialyltransferases gene expression was extracted from normalized counts with minimum 10 transcript counts across the conditions the dataset for in HT29-MTX WT and ΔMUC1 cells. Data were presented as normalized counts and analyzed for statistical significance using Prism GraphPad.

### Cell culture, enzymatic treatments, and confocal microscopy

The human gastrointestinal epithelial cell lines HT29-MTX and HT29-MTX-ΔMUC1 [9], were cultured in 25 cm^2^ or 75 cm^2^ flasks in Dulbecco’s modified Eagle’s medium (DMEM) containing 10% fetal calf serum (FCS) at 37°C in 5% CO2. For confocal microscopy, HT29-MTX and HT29-MTX ΔMUC1 cells were seeded on coverslips (8 mm diameter, #1.5) in 24-well plates to reach confluency at day 3 and grown for a total of 7 days. For lectin-based on-cell ELISA, HT29-MTX WT and ΔMUC1 cells were grown in 96 well plates to reach confluency at day 3 and grown for a total of 7 days. For bacterial infection assays, HT29-MTX WT and ΔMUC1 cells were seeded in 12-well plates to reach confluency at day 3 and grown for a total of 7 days. For neuraminidase treatment, cultures were washed with DPBS and incubated with 100 U/mL α2,3/6/8/9 neuraminidase A (P0722L, New England BioLabs) or 200 U/mL α2,3 neuraminidase S (P0743L, New England BioLabs) in DMEM w/o FCS and incubated for 3 hours at 37°C temp. Afterwards, cells were washed two additional times with DPBS. Lectin staining was performed by incubation with 1:200 Sambucus Nigra Lectin (SNA, 2 mg/mL, B-1305-2, Vector Laboratories) or 1:100 Maackia Amurensis Lectin II (MALII, 1 mg/mL, B-1265-1, Vector Laboratories) in DMEM with 10% FCS for 30 minutes on ice followed by two washes with cold DPBS. Cells were fixed with 4% paraformaldehyde in DPBS (Thermo Fisher) for 30 minutes at RT. Fixation was stopped by adding 50 mM NH₄Cl in DPBS for 15 minutes. Cells were washed twice and incubated in binding buffer (0.2% BSA, Sigma-Aldrich in DPBS) for 30 minutes. Coverslips were incubated with the α-SiiE polyclonal rabbit serum or α-MUC1 antibody 214D4 (in-house) at 1:100 dilution in binding buffer for 1 hour, washed three times with binding buffer, and incubated with Alexa Fluor-647-conjugated α-rabbit IgG secondary antibody (1:200, A21245, Thermo Fisher) Alexa Fluor-568-conjugated α-mouse IgG secondary antibody (1:200, A11029, Thermo Fisher), Alexa Fluor-488-conjugated α-biotin secondary antibody (A6374, Thermo Fisher), and DAPI at 2 μg/mL (D21490, Invitrogen) for 1 hour. Coverslips were washed three times with DPBS, rinsed in Milli-Q water, dried, and embedded in Prolong Diamond Antifade Mounting Solution (Thermo Fisher) and allowed to harden. Images were collected on a spinning disk Olympus SpinSR10 system equipped with a Yokogawa W1-SoRa spinning disk mounted on a IX83 stand with an ORCA Flash 4.0 camera (Olympus, Leiderdorp, the Netherlands). The system was run in confocal mode using 40x oil objective (UPLXAPO, NA1.4). Z slices imaged at 40x were 0.23 µm. Fluorescence was recorded upon sequential excitation by the lasers (Coherent, OBIS) with appropriate emission filters and exposure time. The main dichroic mirror was a quadband (D405/488/561/640 nm). Orthogonal views were displayed in cellSens software (Olympus).

### Lectin-based on-cell ELISA

7-day HT29-MTX WT and ΔMUC1 cultures in 96-well plates were washed with DPBS and 50 μL of either FCS-free DMEM, followed by treatment with neuraminidases as described above. Cells were washed with 100 μL DMEM with 10% FCS, transferred to ice, and 5 μg/mL MALII (Vector Laboratories) or 5 μg/mL SNA (Vector Laboratories) in 50 μL DMEM with 10% FCS (cold) was added and incubated for 30 minutes on ice. Afterwards, wells were washed twice with DPBS and fixed for 30 minutes with 100 μL 4% PFA in DPBS (15670799, Thermo Scientific). Cells were washed twice with DPBS and quenched with 100 μL of 50 mM NH₄Cl in DPBS for 10 minutes. Cells were blocked with 100 μL 0.2% BSA in DPBS for 30 minutes, after which 50 μL of secondary antibody (anti-streptavidin-HRP, 1:1000, 1 μg/mL, Sigma-Aldrich) in binding buffer was added for 1 hour. Cells were washed three times with DPBS, and 50 μL of TMB substrate (3,3’,5,5’-Tetramethylbenzidine, 34028, Thermo Fisher) was added and incubated for 10 minutes until a color shift appeared. The reaction was stopped with 50 μL of 2N H₂SO₄. Absorbance was measured at 450 nm. Absorbance of the secondary antibody control well, and blank were subtracted from the sample absorbance to obtain the values.

### Bacterial invasion assays

Overnight *Salmonella* cultures were diluted 1:1000 in 25 mL LB in a 250 mL Erlenmeyer flask and incubated until OD_600_=1 was reached. OD_600_ was adjusted to 0.24, cells were spun down at 8000 × g for 2 minutes and resuspended in DPBS. For the lectin blockage assay, 7-day HT29-MTX WT and ΔMUC1 cultures in 12-well plates were washed once with DPBS. Lectins Sambucus Nigra Lectin (SNA, biotinylated, Vector Laboratories) and (MALII, Vector Laboratories) were diluted to a concentration of 12 μg/mL in DPBS, and 400 μL was added to each well and incubated for 30 minutes at 37°C. Afterwards, cells were washed twice with DPBS, and 1 mL DMEM without FCS was added. Neuraminidase treatments were performed as described above followed by two washes with DPBS, followed addition of 1 mL DMEM without FCS. 30 μL of bacterial suspension was added which represents a multiplicity of infection (MOI) of 15 and incubated for 1 hour at 37°C. Infected cultures were washed twice with 1 mL DPBS, and DMEM with 300 μg/mL Gentamycin (G1914, Sigma-Aldrich) was added to each well and incubated for 1 hour at 37°C to kill extracellular bacteria. Cells were then washed twice with DPBS and lysed with lysis buffer (0.1% Triton X-100 in DPBS, Sigma-Aldrich) for 10 minutes at 37°C. Serial dilutions were made and plated on LB agar plates. The next day, colonies were counted.

### MUC1 immunoprecipitation

HT29-MTX and HT29-MTX ΔMUC1 cells were seeded on 10 cm² culture dishes to reach confluency at day 3 and grown for a total of 7 days. For harvest, cells were washed once with 1 mL of ice-cold DPBS and collected by scraping in 1 mL of cold DPBS, spun down, supernatant removed, and pellets frozen until use. HT29-MTX WT and ΔMUC1 cell pellets were lysed in 1.5 mL of lysis buffer (25 mM Tris-HCl pH 7.5; 150 mM NaCl; 0.5 mM EDTA; 1% NP40) supplemented with 1× HALT protease inhibitor cocktail (78429, Thermo Fisher Scientific). After adding lysis buffer, samples were put on ice and extensively pipetted every 10 minutes for 1 hour. Tubes were incubated with rotation overnight at 4°C to continue lysis. Lysates were centrifuged at 17,000 × g for 10 minutes at 4°C to remove debris. Protein concentrations were determined using a BCA Protein Assay Kit (23225, Thermo Fisher Scientific). Total protein concentrations were normalized across samples using lysis buffer. Pre-capture lysate samples were stored for analysis. Streptavidin agarose beads (20353, Thermo Fisher Scientific) were equilibrated with wash buffer (25 mM Tris-HCl, pH 7.5; 150 mM NaCl; 0.5 mM EDTA; 0.1% NP40) and incubated with biotinylated anti-MUC1 214D4 antibody. The 214D4 antibody (a kind gift of Dr. John Hilkens, NKI, The Netherlands) was biotinylated using the EZ-Link^TM^ Sulfo NHS-SS Biotinylation Kit (21445, Thermo Fisher Scientific) according to the manufacturer’s instructions, adding a biotin and cleavable linker to the antibody. 40 μg of biotinylated 214D4 was added per 40 μL of beads-slurry and rotated at 4°C for 2 hours. Beads were centrifuged at 2000 × g for 3 minutes and washed with cold DPBS before resuspension in 1.5mllysate. The bead–lysate mixtures were incubated overnight at 4°C while rotating. After binding, beads were centrifuged and supernatants were collected as after-capture samples. Beads were washed twice with DPBS. Elution was performed by incubating the bead pellet with 200 μL of 50 mM DTT in DPBS at 50°C for 1 hour to cleave the NHS-SS linker, followed by centrifugation at 2000 × g. Eluates were collected and 12.5 μL 5× Laemmli buffer was added to 50 μL of eluate. Beads were boiled in 1.25× Laemmli buffer for 10 minutes at 95°C, centrifuged, and supernatants collected. 40 μg of original lysate and 10 μL of after-capture, elution and boiled beads fractions were loaded on gel for Western blot or Coomassie staining.

### MUC1 immunoblots

A mucin-optimized SDS-PAGE gel was prepared as previously described [32]. 30 µg of protein was loaded into each well and run in TAE buffer with 0.1% SDS at 80 mA for 2.5 hours. Proteins were transferred to a PVDF membrane using wet transfer for 3 hours at 90 V at 4°C in transfer buffer (25 mM Tris, 192 mM glycine, Merck; 20% methanol, Merck). Membranes were blocked with 5% BSA in TSMT buffer (20 mM Tris, 150 mM NaCl, Merck; 1 mM CaCl₂, Sigma-Aldrich; 2 mM MgCl₂, Merck; adjusted to pH 7 with HCl, and 0.1% Tween 20, Sigma-Aldrich) for 1 hour at room temperature (RT). Afterwards, membranes were washed with TSMT and incubated with 214D4 (1:2000, MUB2006P, Nordic-MUbio) in TSMT containing 1% BSA for 1 hour at RT. Membranes were washed again with TSMT and incubated with an α-mouse IgG secondary antibody (A2314, Sigma-Aldrich) diluted 1:8000 in TSMT with 1% BSA for 1 hour at RT, followed by washes with TSMT and TSM. Blots were developed using the Clarity Western ECL substrate kit (Bio-Rad) and imaged with a Gel-Doc system (Bio-Rad). Quantification was performed using ImageJ software.

### MUC1 glycan arrays

MUC1 glycopeptide microarray from ZBiotech (MUC1 Glycopeptide Array2, 10619-16S) includes a library of 124 glycopeptides on the same MUC1 VNTR repeat sequence of 23 amino acids and one non-glycosylated control peptide. The peptides are modified with one or more mucin-type O-glycan structures including core 2 (Gal(β1-3)[GlcNAc(β1-6)]GalNAc(α1-*O*)Ser/Thr), α2,3-sialylated TF antigen (α2-3-sTF: Neu5Ac(α2-3)Gal(β1-3)GalNAc(α1-*O*)Ser/Thr), α2,6-sialylated TF antigen (α2,6-sTF: Neu5Ac(α2-6)Gal(β1-3)GalNAc(α1-*O*)Ser/Thr), and disialylated TF antigen (di-sTF: Neu5Ac(α2-3)Gal(β1-3)[Neu5Ac(α2-6)]GalNAc(α1-*O*)Ser/Thr). The O-glycans groups are attached to the peptide backbone with varying spacing and density (Fig. 2G). For incubation with the MUC1 glycopeptide array, SiiE was harvested from 500 ml bacterial supernatant of an overnight culture by concentration using a 100 kDa cut off spin column to 500 µl, containing 2.1 µg/µL total protein. The supernatant of a Δ*siiE* culture supernatant was prepared as negative control (1.7 µg/µL total protein). SiiE samples were diluted in binding buffer (1% BSA, Sigma-Aldrich A8577, in 20 mM Tris-HCl pH8 with 10 mM CaCl₂) to final concentrations of 100 ng/µL (low) and 300 ng/µL (high). 200 µL of samples SiiE low and high, ΔSiiE (50 ng/µL, negative control) and PNA (5 mg/ml, L10707, Vector Laboratories) (5 µg/mL, quality control) were incubated with the MUC1 Glycopeptide Array2 for 1.5 hours at 20°C. The array was then washed three times with wash buffer (20 mM Tris-HCl, pH 8.0, supplemented with 10 mM CaCl₂ and 0.05% Tween-20) 1 minute each at 20 °C. Subsequently, the array was incubated with 200 uL of polyclonal anti-SiiE antibody at 10ug/mL (Ab1; α-SiiE rabbit serum provided by Prof. Dr. Michael Hensel, Universität Osnabrück, Germany) for 1 hour, then with 200 uL of biotinylated anti-rabbit IgG antibody at 10 ug/mL (Sigma-Aldrich B8895; Ab2) for 1 hour, and finally with 200 uL of AlexaFluor-647-labeled streptavidin at 1 μg/ml (Molecular Probes) for fluorescence detection. All incubation steps were performed in 200 µL of 20 mM Tris-HCl buffer (pH 8.0) containing 10 mM CaCl₂, with three washing steps between each incubation. Imaging and data analysis were performed as previously described [42]. Fluorescence parameters were optimized to balance signal-to-noise ratio and avoid saturation across experiments. Results given are plotted as an average of three replicates for binding signals. The negative control arrays that were incubated with Δ*siiE* sample or with anti-SiiE/anti-rabbit antibodies in the absence of SiiE sample did not produce detectable binding signals.

**Figure 2.**
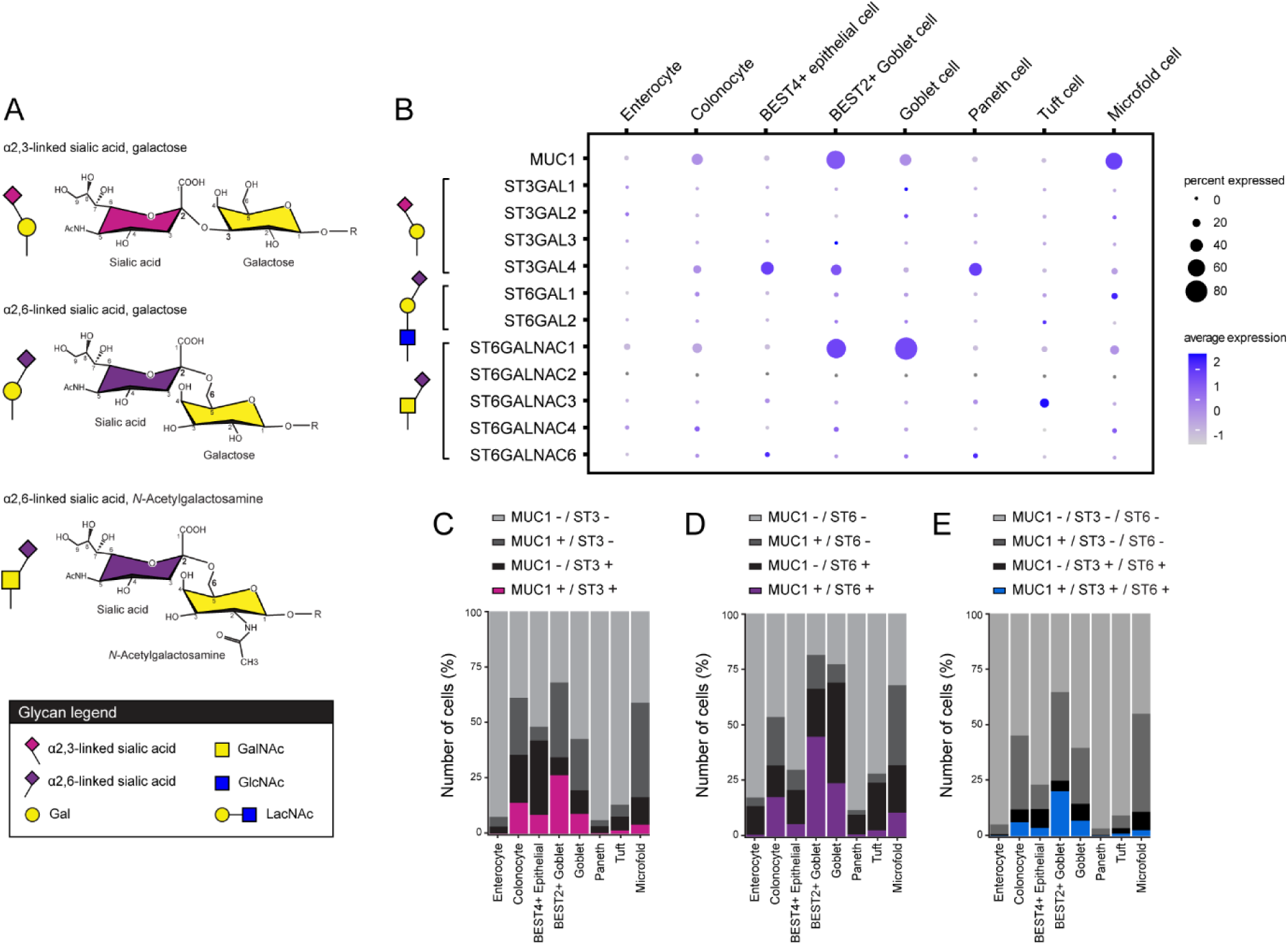
Single cell RNAseq analysis of MUC1 and sialyltransferase expression in intestinal epithelial cell types. A) Schematic representations and molecular structures of α2,3-linked sialic acid to galactose, and α2,6-linked sialic acid to GalNAc or galactose. Glycan representations are according to Symbol Nomenclature for Glycans (SNFG) with distinguishing colors for α2,3 and α2,6 linkages. B) Single-cell RNA-sequencing analysis of MUC1 and sialyltransferease expression in adult donors from the Gut Cell Atlas database. Expression levels and percentages of MUC1, α2,3-linkage specific ST3GAL (ST3), α2,6-linkage specific ST6GAL and ST6GALNAC (ST6) sialyltransferase genes in intestinal epithelial enterocytes, colonocytes, BEST4+ epithelial cells, BEST2+ goblet cells, Paneth cells, tuft cells, and microfold cells are indicated. C) Stacked bar graphs depicting percentages of cells expressing MUC1, ST3, or co-expressing both. D) Stacked bar graphs depicting percentages of cells expressing MUC1, ST6GAL/ST6GALNAC, or co-expressing both. E) Stacked bar graphs depicting percentages of cells expressing MUC1, ST3 and ST6, or co-expressing all.

### O-glycan release

HT29-MTX and HT29-MTX ΔMUC1 cells were seeded on 10 cm² culture dishes to reach 100% confluency at day 3 and grown for a total of 7 days. For neuraminidase treatment, cultures were washed with DPBS and incubated with 200 U/mL α2,3 neuraminidase S (P0743L, New England BioLabs) in DMEM w/o FCS and incubated for 3 hours at 37°C temp. Afterwards, cells were washed two additional times with ice-cold DPBS. For harvest, cells were washed once with 1 mL of ice-cold DPBS and collected by scraping in 1 mL of cold DPBS, spun down, supernatant removed, and pellets frozen at -80°C until use. For lysis, the cell pellets were resuspended in lysis buffer (50 mM Tris HCl, 100 mM NaCl) with approximately 20,000 cells/µL or 2 µg/µL protein. The cell pellets were agitated by pipetting until no agglomerated cells were observed, followed by 30 min sonication and heating for an additional 30 min at 60°C with shaking. Protein lysate or MUC1 IP sample (25 µL) was loaded on PVDF membranes (MultiScreenHTS IP Filter Plate, 0.45 μm, Millipore) and allowed to bind as described previously [43]. A de-N-glycosylation was performed on the proteins attached to the membrane followed by a nonreductive O-glycan release using 25 μL of release agent (21% hydroxylamine, 17% DBU and 0.3 nM maltopentaoase (DP5) in water) and incubated in a moisture box at 37 °C for 75 min. The liquid containing the released O-glycan products was recovered from the membrane by centrifugation and added to 1 mL of 100% ACN containing 2 mg of magnetic hydrazide beads (MagSi-S Hydrazide beads 1 μm). After two washes with ACN, the O-glycans were eluted from the hydrazide beads in 50 μL of 2-AB reagent (500 mM 2-AB, 116 mM PB in 45:45:10 (%, v/v) methanol:water:acetic acid). The 2-AB labeling reaction was performed by incubation for 2.5 h at 50 °C, purified by cotton HILIC SPE (loading step: 99% ACN in water, and washes with 100% ACN) and 2-AB labeled O-glycans were eluted in water, following further purification by PGC SPE as described previously [43]. The samples were dried and reconstituted in 20 μL water for C18 nano-scale Liquid Chromatography (nano-LC) coupled with electrospray ionization (ESI) mass spectrometry (MS) analysis.

### Liquid Chromatography–Mass Spectrometry of labeled O-glycans

For LC-MS analysis, 2 μL per sample (10% of total) was injected per analysis and all samples were analyzed at least 3 times. The glycans were separated by an EASY-nLC 1,200 UHPLC (Thermo Fisher Scientific, Germering, Germany), mobile phase A consisted of 0.1% formic acid in water and using a gradient from 2% to 32% mobile phase B in 20 min (0.1% formic acid/80% ACN in water). A single analytical column setup (Reprosil-Pure-AQ C18 phase, Dr. Maisch, 1.9 μm in particle size, ∼25 cm in column length) with an emitter was interfaced to an Orbitrap Fusion Lumos MS (Thermo Fisher Scientific, San Jose, USA) via a nanoSpray Flex ion source. A precursor MS scan (*m/z* 275–1,700, positive polarity) was acquired in the Orbitrap at a nominal resolution of 120,000, followed by Orbitrap higher-energy C-trap dissociation (HCD)-MS/MS at a nominal resolution of 30,000 of the 10 most abundant precursors in the MS spectrum (charge states 1 to 4). A minimum MS signal threshold of 50,000 was used to trigger data-dependent fragmentation events. HCD was performed with an energy of 27% ± 5%, applying a 10 s dynamic exclusion window. In addition to the HCD event, an ion-triggered fragmentation event was initiated using CID with an energy of 32% on precursors of which the HCD spectrum contained one of the following *m/z* values; 204.0866 (HexNAc), 342.166 (HexNAc-2AB), 285.144 (Fuc-2AB) or 301.1394 (Hex-2AB).

### Data Analysis

MS1 feature detection in the raw files was performed using the Minora Feature Detector node in Thermo Proteome Discoverer 2.5.0.400 (Thermo Fisher Scientific Inc.). The [M + H] values of the resulting features were imported into GlycoWorkbench 2.1 (build 146) and matched to glycan compositions with 0 to 2 pentoses, 0 to 10 hexoses, 0 to 10 *N*-acetylhexosamines, 0 to 3 fucoses, 0 to 4 *N*-acetylneuraminic acids, 0 to 2 *N*-glycolylneuraminic acids, and a 2-AB label. The complete list of identified compositions was imported into Skyline 24.1.0.414 (ProteoWizard) using the Molecule Interface. Extracted ion chromatograms were generated for the first three isotopologues of each glycan, chromatographic peaks were selected based on accurate mass (>−1 ppm, <1 ppm), isotopic dot product (idotp; > 0.80) and a signal intensity threshold of 2× 10^5^ in the extracted total MS1 spectrum. Following these curation steps, each chromatographic peak had to be detected in at least one of the 3 replicates. Finally, total area normalization was performed for the subset of O-GalNAc glycans to obtain the relative abundances per glycan in each sample. Glycan structural assignments were deduced (manually) from MS/MS patterns, retention behavior in LC-MS, and biosynthetic knowledge of glycan pathways. Glycan cartoons are presented as structural representations based on this combined evidence and established biosynthetic logic. For assignment of O-glycans containing α2,3-linked *N*-acetylneuraminic acid, the O-glycan repertoire of untreated samples and cells treated with α2,3-specific Neuraminidase S were compared.

### Human InTESTine^TM^ *ex vivo* intestinal cultures and infections

Williams E (WE) buffer was prepared as previously described [44]. Human intestinal ileum and colon tissue was obtained from a human adult patient undergoing laparoscopic right hemicolectomy surgery for colon carcinoma removal. Ethical approval for the use of human intestinal tissue was obtained from the hospital board. Because samples were collected once and anonymously, they are not subjected to the “Wet medisch-wetenschappelijk onderzoek met mensen” (WMO) in the Netherlands. Prior to surgery, informed consent was obtained from the patient. Directly after dissection of the intestinal tissue, the tissue was transported to the pathologist and healthy considered segments for the distal ileum and transverse colon were donated for research purposes. For transport and handling, tissues were placed in ice-cold WE buffer supplemented with 1% penicillin/streptomycin. InTESTine™ experiments were generally performed as previously described for a 24-well plate [44, 45] with a few modifications. In brief, a puncher (BAHCO PONS, B400.010, 10 mm) was used to generate 0.246 cm^2^ punches from the distal ileum and transverse colon segments followed by mounting in InTESTine™ inserts within 2 hours after surgical removal. 125 μL cold WE buffer without antibiotics was added to the apical side and the setup placed in a humidified incubator at 37°C with 95% O_2_, 5% CO_2_ for pre-incubation and to slowly warm-up to 37 °C for a period of 30 min (t = -30 min). At t=0 h, inserts were transferred to a new 4-well plate pre-filled with 875 μL WE without antibiotics + 4% BSA in the basolateral compartment that was pre-warmed at 37 °C. 62.5 μL of the apical solution was carefully removed to avoid disturbing the apical mucus layer and replaced by dosing solution consisting of WE without antibiotics without bacteria (blank) or with *Salmonella* Enteritidis WT (both at 2x10^8^ CFU/mL). Final apical concentrations were 1*10^8^ CFU/mL for bacteria. All apical doses were pre-warmed to 37°C for 30 minutes before addition to the tissue. Next, the setup was incubated for 1 h at 37°C with 95% O_2_, 5% CO_2_. Apical and basolateral samples were harvested and stored for analysis. To assess the viability of the *ex vivo* intestinal segments, the cytosolic enzyme lactate dehydrogenase (LDH) was measured in the apical and basolateral supernatants and compared to the LDH level in homogenized tissue segments (t=0 h), using an LDH kit (Sigma-Aldrich, 4744926001) as previously described [44, 45]. To determine bacteria associated with the tissue, intestinal segments were dismounted and washed three times with sterile PBS (Fisher Scientific, 10010056) and homogenized in 1 mL sterile WE medium without antibiotics using the Bead Mill Homogenisator (VWR). Colony-forming units were quantified by plating 10 µL of serial 10-fold dilutions of apical samples, homogenized segments, and basolateral samples in duplicate on square TSA plates containing a 6x6 grid using the drop plating method and incubated overnight at 37°C. The next day, bacterial colonies were counted. For microscopy, intestinal segments were fixed in the InTESTine™ insert with 10% formalin (VWR, FOR0150AF59001; 150 μL apical, 875 μL basolateral). Formalin-fixated inserts were kept in fixative for 24 hours after which the fixative was replaced by 70% ethanol and samples were stored at RT until further processing.

### Tissue paraffin embedding and preparation for immunostaining

Tissues were fixed for 24–48 h in 4% phosphate-buffered formaldehyde, pH 7.0 (ROTI Histofix, Carl Roth). Fixed tissues were placed in embedding cassettes and processed overnight using a HistoCore PEGASUS tissue processor (Leica Biosystems). Samples were dehydrated through a graded ethanol series consisting of four changes of 70% ethanol, two changes of 96% ethanol, and three changes of 100% ethanol, each for 60 min at room temperature. Samples were then cleared in xylene for 60 min at room temperature followed by 60 min at 40°C. Finally, tissues were infiltrated with paraffin in three consecutive 60-min steps at 62°C. Processed tissues were embedded in paraffin using metal embedding mounds (Carl Roth, TT30.1). Paraffin tissue sections of 3-4 μm were cut using a CUT5062 microtome (SLEE medical GmbH, Germany) and mounted onto poly-lysine coated glass slides (0.01%, P4707, Sigma-Aldrich). Slices were air-dried for 30 min and incubated at 60 °C for 2 hours to bind the tissues to the glass surface. Tissues were deparaffinized in 100% xylene followed by a step-wise rehydration from 100% alcohol to 70% to water. A hydrophobic boundary was drawn around tissues on each slide using a PAP pen (Biotium company, #22005), after which slides were placed in a humidity container and covered with either Dulbecco’s PBS without calcium or magnesium (DPBS; D8537, Sigma-Aldrich) for direct immunostaining or hybridization buffer (1.5M NaCl, 1M Tris, 0.1% v/v SDS, 20% formamide, pH 7.4) for FISH/immunostaining.

### Immunostaining of deparaffinized tissue sections

Slides were blocked in blocking buffer (2% BSA, 0.3% Triton X100 in DPBS-/-) for one hour at RT. Staining with primary antibodies and lectins was performed in staining buffer (2% BSA, 0.1% Triton X100 in DPBS-/-) for 1 h at RT. Antibodies and lectins used were α-MUC1 antibody (clone 214D4, 1:100), biotinylated MAL-II (MALII, 1 mg/mL, B-1265-1, Vector Laboratories, 1:100), biotinylated SNA (SNA, 2 mg/mL, B-1305-2, Vector Laboratories, 1:100), Cy5-conjugated SNA (CL-1305, Vector laboratories; 1:100). Slides were washed 3 times 5 min in DPBS-/- and incubated for 30 min in staining buffer containing Alexa Fluor-568-conjugated α-mouse IgG secondary antibody (A11029, Thermo Fisher; 1:200), Oregon Green-488-conjugated NeutrAvidin (A6374, Invitrogen; 1:200) and DAPI (D21490, Invitrogen; 1:500). Slides were washed three times with DPBS-/-, rinsed in Milli-Q water, dried, and embedded in Prolong Diamond Antifade Mounting Solution (Thermo Fisher) and allowed to harden. Images were collected on a Nikon A1 confocal laser microscope equipped with a spectral 32 PMT detector array using a 20x air objective (NA 0.75, CFI PlanFluor multi-immersion) controlled by NIS elements software (NIKON) with default settings to detect DAPI, Oregon Green 488, Alexa Fluor 568 and Cy5.

## Results

### Expression and secretion of the *Salmonella* SiiE giant adhesin under different growth conditions

To identify the optimal growth conditions for SiiE expression by the *Salmonella* Enteritidis strain, we selected four conditions previously reported to result in the highest levels of *siiE* mRNA expression [34]. These conditions included aerobic growth to early and late exponential phase (EEP/LEP, OD_600_ 1 and 2), anaerobic growth until OD_600_ 0.3, and anaerobic growth followed by an oxygen shock. The oxygen shock condition mimics the transition from the anaerobic lumen of the intestine to the more oxygen-rich environment at the intestinal epithelium. *Salmonella enterica* serovar Enteritidis cultures were grown overnight (O/N) and EEP, LEP, anaerobic and oxygen shock cultures were prepared. Expression of SiiE protein on the bacteria was visualized using immunofluorescence and quantified using ImageJ (Fig. 1B,C). In the overnight culture, SiiE expression was relatively low with positive staining of around 1% of bacteria. SiiE expression was greatly induced in most other conditions and highest in the EEP, followed by anaerobic growth, oxygen shock and LEP conditions. Compared to aerobic growth, SiiE expression was relatively high under anaerobic conditions and significantly induced by oxygen shock after anaerobic growth.

While SiiE is secreted via the bacterial T1SS, the protein can remain associated with the bacterial surface [32]. To quantify the cell wall associated and released SiiE under the different conditions, bacterial pellets and concentrated supernatants were analyzed by immunoblot. In the bacterial pellets, we observed multiple SiiE-reactive bands ranging between 250 and 600 kDa, while the supernatant fractions contained a single band corresponding to the predicted full-length SiiE molecular weight of 600 kDa (Fig. 1D, E). Quantification of the bands in the pellet fraction demonstrated that SiiE expression was significantly higher in the EEP and LEP conditions compared to the O/N condition (Fig. 1E). Under anaerobic conditions and oxygen shock, expression levels were relatively low compared to the aerobic conditions. In line with the immunofluorescence analysis, a significant induction of SiiE expression was observed after oxygen shock (Fig. 1F). In the secreted fractions, SiiE expression was high in the O/N fraction and of comparable intensity in the EEP and LEP growth conditions. SiiE was not detectable in the secreted fraction of the anaerobic growth and oxygen shock conditions. In conclusion, SiiE expression on the bacterial surface was highest under aerobic conditions, particularly during the early exponential phase. Secreted SiiE was detectable in high quantities under both exponential growth and overnight conditions. Oxygen shock, a condition that occurs when bacteria reach the intestinal epithelium where the receptor MUC1 is expressed, significantly induced SiiE expression on the bacterial surface.

### ST3 and ST6 transferases are co-expressed in MUC1-positive intestinal epithelial cells

We previously demonstrated that the interaction of SiiE with MUC1 is dependent on sialic acids. These residues are commonly found as terminal monosaccharides on O-glycan structures and can occur as α2,3-linked sialic acids attached to a galactose (Gal), as well as α2,6-linked sialic acids attached to either Gal or *N*-acetylgalactosamine (GalNAc). The α2,3- and α2,6-linked sialic acids are generated by of the sialyltransferases ST3GAL, ST6GAL and ST6GALNAC, respectively (Fig. 2A). To explore the potential *in vivo* sialylation patterns of MUC1, we determined the expression of MUC1 and the ST3GAL, ST6GAL and ST6GALNAC sialyltransferases family members in various cell types of the human intestine using the online Gut Cell Atlas single-cell RNAseq platform. Expression and co-expression levels of MUC1 and the ST3GAL, ST6GAL and ST6GALNAC sialyltransferases were analyzed across different intestinal epithelial cell types from healthy adult donors. MUC1 was expressed in colonocytes, goblet cells, BEST2+ goblet cells, goblet cells and microfold cells (Fig. 2B), with markedly lower expression in enterocytes compared to colonocytes, indicating limited expression in the small intestine. Among the α2,3 sialyltransferases, ST3GAL4 showed the highest expression, predominately in colonocytes, BEST4+ epithelial cells, BEST2+ goblet cells and Paneth cells. For the α2,6 sialyltransferases, ST6GALNAC1 presented the highest expression, with presence in colonocytes and particularly high levels in BEST2+ goblet cells and goblet cells. Expression of ST6GAL enzymes, the sialyltransferase members responsible for the α2,6-sialylation of *N-*acetyllactosamine (LacNAc) motifs, was low and only a small percentage of microfold cells expressed ST6GAL1. This suggests that the overall occurrence of α2,6-linked sialic acids on galactoses are relatively scarce on the intestinal epithelium of healthy adults. Next, we assessed co-expression patterns of MUC1 and sialyltransferases across epithelial cell types. Co-expression of MUC1 with either ST3GAL or ST6GAL transferases was observed in colonocytes, BEST4+ epithelial cells, goblet cells and highest in BEST2+ goblet cells where double positive cells amounted to 25-50% of the total cell population (Fig. 2C, D). Triple co-expression of MUC1, ST3GAL and ST6GAL genes was also highest in BEST2+ goblet cells with 20% of the total population (Fig. 2E). These data suggest that MUC1 expression in intestinal epithelial cells can coincide with expression of both ST3GAL and ST6GALNAC transferases and therefore could carry both α2,3-and α2,6-linked sialic acid structures.

### Secreted SiiE interacts with diverse O-glycan structures on MUC1

To investigate the capacity of *Salmonella* SiiE to interact with different glycan groups on MUC1, we used the full length SiiE containing 53 lectin-like domains in combination with a MUC1 glycan array comprising 124 glycopeptides. Each glycopeptide consisted of a single 23 amino acid MUC1 VNTR domain with one or more O-glycans on serine and threonine residues, including both sialylated and non-sialylated forms, with varying spacing and density. Secreted full-length SiiE was purified from bacterial supernatant of stationary cultures (OD_600_=2.0) resulting in a relatively pure fraction (Fig. 3A). The purified SiiE and the lectin PNA that binds to exposed Gal of type 3 (Galβ1,3-GalNAc-) motifs were incubated with the MUC1 glycan array, and fluorescence intensity was measured to quantify interaction. As expected, PNA binding was observed on MUC1 glycopeptides presenting a terminal Gal residue in the absence of sialic acid capping (Fig. 3B). SiiE displayed a more diverse binding pattern as it interacted with all classes of MUC1 glycopeptides (Fig. 3B). Strongest SiiE binding was observed to MUC1 with a core 2 O-glycan, confirming previous reports of its recognition of GlcNAc residues [9, 33]. Moderate binding to glycopeptides containing the α2,3-linked sialic acid (α2,3-sTF) on core 1 was observed. This binding was highly dependent on position of the glycan group and seemed to decrease with increased glycan density. SiiE also interacted with glycopeptides containing α2,6-linked sialic acid (α2,6-sTF) including some peptides with high glycan density. Moderate binding to disialylated peptides containing both an α2,3-linked and α2,6-linked sialic acid (di-sTF) was also observed with a preference for lower glycan density peptides. We conclude that SiiE in its secreted form is able to interact with diverse glycan structures and glycan motifs on MUC1 peptides. Interestingly, positioning of the glycan structure groups on the MUC1 peptide determined specificity and the optimal position was different for glycans containing α2,3-linked sialic acid, α2,6-linked sialic acid or both.

**Figure 3.**
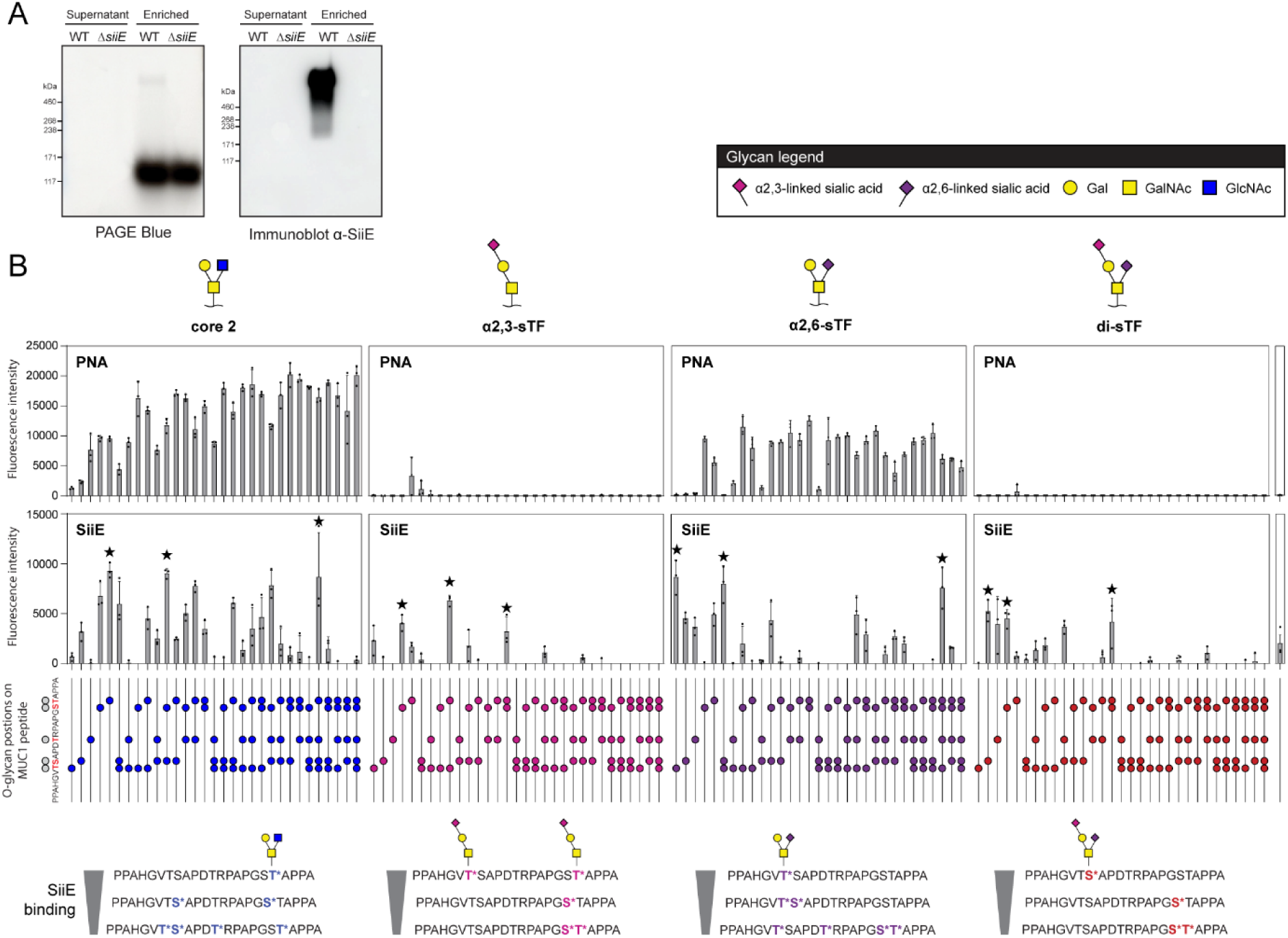
SiiE interacts with diverse glycosylated MUC1 peptides in a MUC1 glycan array. A) PAGE blue and immunoblot analysis of *Salmonella enterica* serovar Enteritidis supernatant and concentrated supernatant fractions demonstrating high SiiE concentrations in the concentrated supernatant fraction of WT bacteria. B) Interaction of SiiE and galactose-binding lectin PNA with diverse O-glycosylated MUC1 peptides in a MUC1 glycan array. The array consists of the MUC1 repeat domain peptide decorated with non-sialylated core-2 glycans (core 2) at different serine and threonine residues, and peptides decorated with α2,3-sialylated core-1 glycans (α2,3-sTF), α2,6-sialylated core-1 glycans (α2,6-sTF), α2,3 and α2,6-sialylated core-1 glycans (di-sTF) and a non-glycosylated peptide control. The position of the glycan modification on the MUC1 peptide is indicated by a colored circle. The array was incubated with 5ng/µl PNA to detect exposed galactose residues or 300 ng/µl enriched SiiE fraction. For each glycan category, the three MUC1 glycopeptides with strongest SiiE binding are indicated by a star and depicted in the bottom. Graphs represent mean ± SD from three independent biological replicates.

### Colonic HT29-MTX cells express MUC1 and α2,3- and α2,6-linked sialic acids on the apical surface

To investigate the presence of α2,3- and α2,6-linked sialic acids on the intestinal epithelium, we utilized the colonic mucus-producing HT29-MTX cell line that we previously used for *Salmonella* invasion studies [9]. Transcriptomics analysis of HT29-MTX WT and ΔMUC1 cultures demonstrated that ST3GAL1-4 were highly expressed in both cell lines and ST3GAL5 and ST3GAL6 were not detectable (Fig. 4A) [46]. For the ST6 sialyltransferases, both ST6GAL1 and ST6GALNAC1 were highly expressed, and ST6GALNAC1 expression was significantly lower in the WT compared to the ΔMUC1 cultures. Next, we determined the presence of α2,3- and α2,6-linked sialic acids on the surface of confluent HT29-MTX WT and ΔMUC1 monolayers by staining with lectins MALII to detect α2,3-linked sialic acids and SNA to detect α2,6-linked sialic acids. HT29-MTX cultures were left untreated or treated with a broad-spectrum neuraminidase that removes α2,3/6/8/9 sialic acid linkages or with an α2,3-specific neuraminidase to selectively cleave α2,3-linked sialic acids. In untreated cultures, MALII staining showed a strong signal at the apical surface of HT29-MTX WT cells, suggesting a high abundance of α2,3-linked sialic acids. Additionally, MUC1 was highly expressed on the apical surface, with regions of co-localization observed between MALII and MUC1 (Fig. 4B). After treatment with either the broad-spectrum or α2,3-specific neuraminidase, MALII staining was reduced, indicating effective removal of α2,3-linked sialic acids by both enzymes (Fig. 4B). SNA lectin staining revealed a clear signal at the apical surface of HT29-MTX WT cells, indicating the presence of α2,6-linked sialic acids. Treatment with the broad-spectrum neuraminidase led to a significant reduction of SNA staining, indicating that α2,6-linked sialic acids were removed (Fig. 4C). Under control conditions, no apparent co-localization of SNA with MUC1 was observed. However, after treatment with the α2,3-specific neuraminidase, SNA staining changed compared to the untreated control, and some co-localization of SNA and MUC1 was observed (Fig. 4C). The results suggested that certain α2,6-sialic acid epitopes were masked by the presence of α2,3-linked sialic acids and became exposed upon the removal of α2,3-linked sialic acids. A lectin-based on-cell ELISA was performed to quantify the presence of α2,3- and α2,6-linked sialic acids on the apical surface. Consistent with the immunofluorescence results, the MALII signal was strongly diminished following treatment with both the broad-spectrum and α2,3-specific neuraminidases (Fig. 4D), whereas the SNA signal decreased only after treatment with the broad-spectrum neuraminidase (Fig. 4E). The results confirmed the differential expression and accessibility of α2,3- and α2,6-linked sialic acids on the apical surface of HT29-MTX cells, where α2,3-linked sialic acids are predominant and frequently co-localize with MUC1.

**Figure 4.**
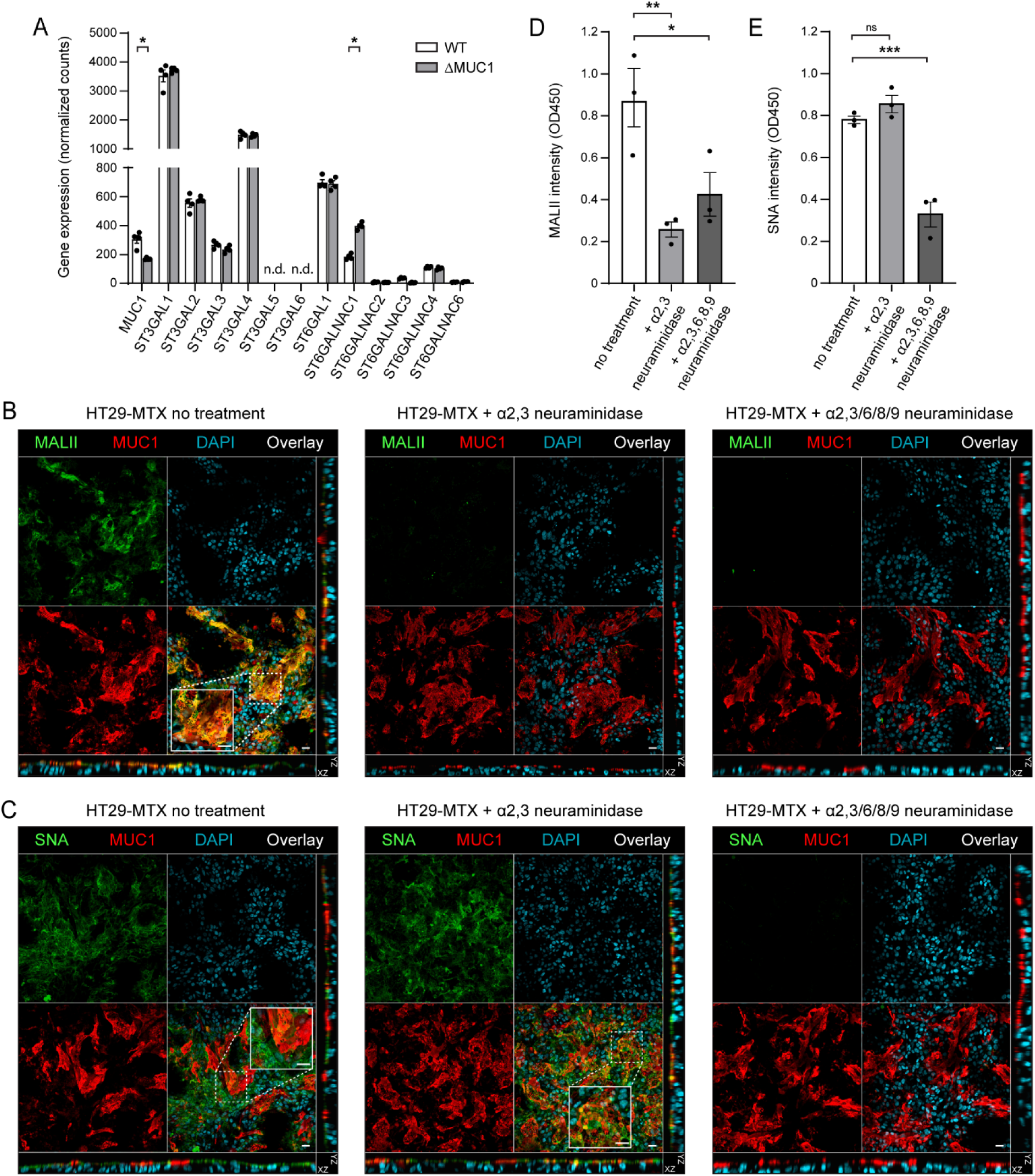
The apical surface of HT29-MTX cells contains both α2,3- and α2,6-linked sialic acids. A) RNAseq expression data of MUC1 and α2,3-linkage specific ST3 and α2,6-linkage specific ST6 sialyltransferase genes in HT29-MTX WT and ΔMUC1 cultures. Statistical test: two-tailed paired t-test. B, C) Immunofluorescence of HT29-MTX WT cultures without treatment or treated with α2,3-specific neuraminidase or α2,3/6/8/9 broad specificity neuraminidase strained for DAPI (blue), MUC1 (red) or MALII/SNA (green). D, E) On-cell ELISA quantification of SNA and MALII binding to HT29-MTX WT cultures without treatment or treated with α2,3-specific neuraminidase or α2,3/6/8/9 broad specificity neuraminidase. Statistical test: one-way ANOVA with Dunnett’s correction comparing WT to all conditions. Graphs represent mean ± SD (A) or ± SEM (B,C) from three independent biological replicates. ns: not significant; * *p*<0.05; **** *p*<0.0001. Scale bars: 20 µm.

### O-glycomics analysis of HT29-MTX cultures and immunoprecipitated MUC1

Next, we performed O-glycomic analysis to identify the specific sialylated O-GalNac glycans present on HT29-MTX cells and MUC1. Total cell lysates were prepared from 7-day HT29-MTX wild type and ΔMUC1 cultures (WT-TCL and ΔMUC1-TCL), as well as from wild type cultures treated with an α2,3-specific neuraminidase. In parallel, 7-day HT29-MTX wild type and ΔMUC1 cultures were used for immunoprecipitation of MUC1 using a biotinylated α-MUC1 antibody and a streptavidin column (MUC1-IP; Fig. 5A, B). MS-based O-glycomic analysis of the samples demonstrated the presence of different α2,3- and α2,6-sialylated O-GalNAc structures in WT-TCL, ΔMUC1-TCL and MUC1-IP samples (Fig. 5C, S1B,C,D). Analysis of HT29-MTX cultures treated with α2,3-specific neuraminidase aided in the assignment of the sialic acid linkages in the detected structures (Fig. S1A). The most abundant sialylated O-GalNAc structures were α2,3-sTF (H1N1S1b, a core 1 glycan with one α2,3-linked sialic acid), di-sTF (H1N1S2, a core 1 glycan with one α2,3- and one α2,6-linked sialic acid) and di-sialyl core 2 (H2N2S2, containing two α2,3-linked sialic acids). Less abundant sialylated O-GalNAc structures included α2,6-sTF (H1N1S1c) and sTn (N1S1), both containing an α2,6-linked sialic acid on the GalNAc residue (Fig. 5C). Overall, the repertoire of O-GalNAc structures in the TCL samples and MUC1-IP sample were comparable. However, α2,3-sTF/H1N1S1b and di-sTF/H1N1S2 core 1 were significantly lower in ΔMUC1-TCL compared to WT-TCL and enriched in the MUC1-IP sample (Fig. 5C). Di-sialyl core 2/H2N2S2, on the other hand, showed a reverse pattern with higher levels in ΔMUC1-TCL and lower levels in the MUC1-IP sample (Fig. 5C). In line with this, the MUC1-IP samples had an overall higher level of core 1 structures, and a lower level of core 2 structures as compared to the TCL samples (Fig. 5D, E). The total percentage of O-glycans decorated with an α2,3-linked sialic acid was lower in the ΔMUC1-TCL sample compared to the WT-TCL and highest in the MUC1-IP sample (Fig. 5F). In contrast, no difference in the α2,6-linked sialic acid on core 1 was observed between the samples (Fig. 5G) Together, these data demonstrate that HT29-MTX cultures have a diverse O-glycan repertoire consisting of core 1 and core 2 glycans and that the MUC1 glycoprotein mostly carries sialylated core 1 structures α2,3-sTF and di-sTF. This glycan profile with prominent α2,3-sialylation on Gal residues, while α2,6-sialylation is restricted to GalNAc residues (Fig5C) is broadly consistent with high expression of ST3GAL1, ST3GAL4 and ST6GAlNAC1 as observed in the HT29-MTX RNAseq data (Fig. 4A). Although ST6GAL1 was highly expressed (Fig. 4A), no α2,6-sialylated Gal residues were detected on the O-glycans (Fig. 5C). This outcome may be the result of competition between ST6GAL1 and ST3GAL glycosyltransferases, which preferentially sialylate available LacNAc acceptor structures, and the limited abundance of suitable core 2 O-glycans that serve as substrates for ST6GAL1. This example illustrates the importance of performing O-glycomics analyses to accompany transcriptomics data.

**Figure 5.**
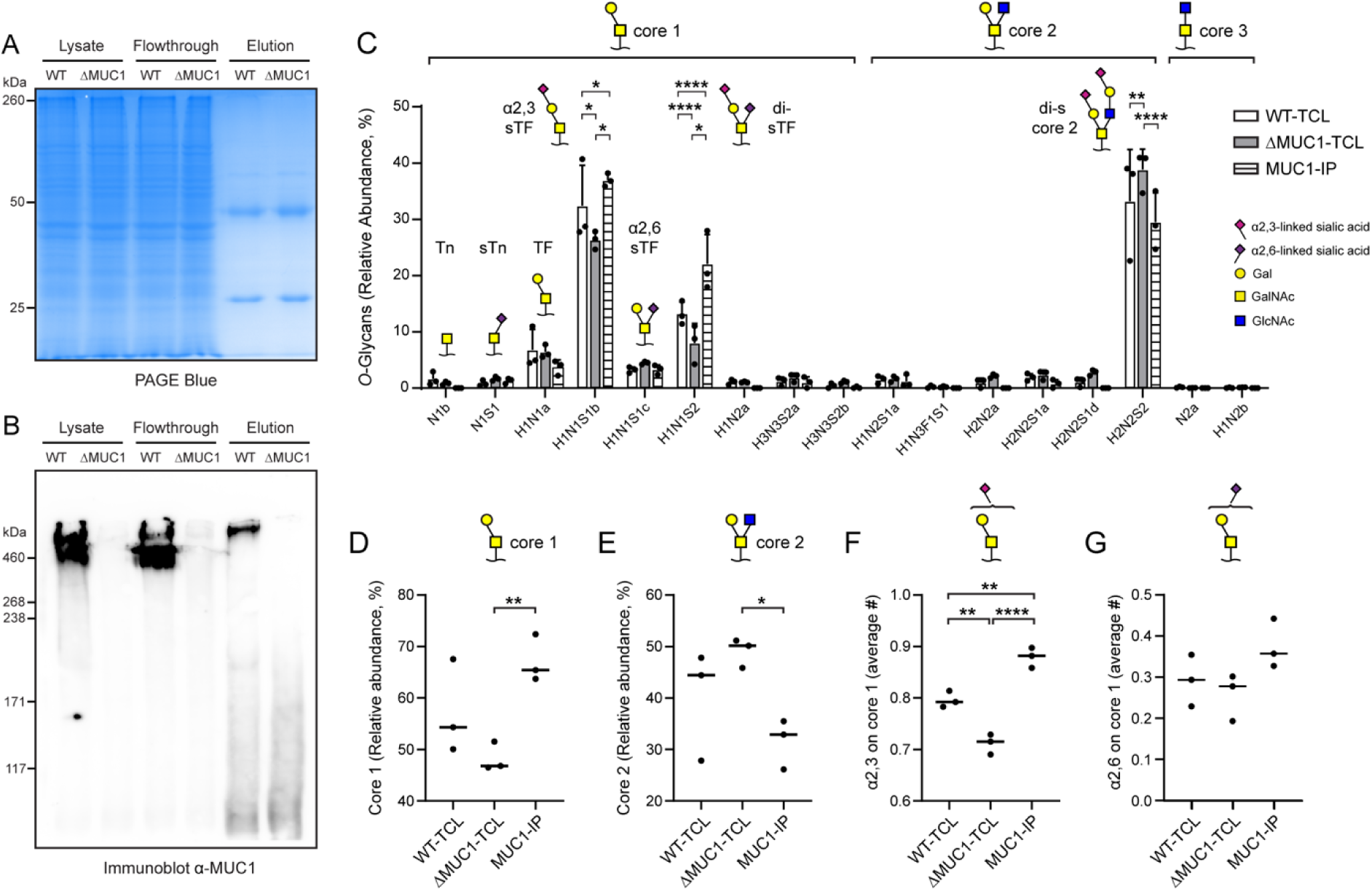
α2,3-sialylated core-1 O-glycan structures are abundant on MUC1 expressed in HT29-MTX cultures. A, B) MUC1 immunoprecipitation (IP) samples from HT29-MTX cultures analyzed by PageBlue and immunoblot analysis. IP fractions include HT29-MTX-WT and HT29-MTX-ΔMUC1 total lysate, flowthrough, and elution. In the PageBlue gel, the light and heavy chains of the eluted anti-MUC1 139H2 antibody are visible at 25 and 50 kDa and the immunoblot demonstrates successful elution of high-molecular weight MUC1 from the WT cultures. C) O-glycomics analysis of O-GalNAc containing structures in HT29-MTX-WT and HT29-MTX-ΔMUC1 total cell lysates (TCL) and MUC1 IP elution (MUC1-IP) samples depicting relative abundance of specific O-glycans. The most abundant detectable structures were TF (no sialic acids), α2,3-sTF (containing a α2,3-linked sialic acid), α2,6-sTF (containing an α2,6-linked sialic acid), di-sTF (containing both an α2,3- and an α2,6-linked sialic acid), di-s core 2 (containing two α2,3-linked sialic acid), Tn (no sialic acids) and sTn (containing a α2,6-linked sialic acid). The schematic representations of the structures with differentially colored α2,3 and α2,6 linkages according to SNFG are included. H = hexose, N = *N*-acetylhexosamine, F = fucose, S = *N*-acetylneuraminic acid. a, b, c and d are added to the glycan name to distinguish LC-separated glycan isomers. The relative total abundance of core 1 structures (D), core 2 structures (E), the average number of α2,3-linked sialic acids on core 1 structures (F) and the average number of α2,6-linked sialic acids core 1 (G) in WT-TCL, ΔMUC1-TCL and MUC1-IP samples was plotted. Graphs represent mean ± SD from three independent biological replicates. Statistical test: two-way ANOVA with Tukey’s correction. ns = not significant; * p<0.05; ** p<0.01; *** p<0.001; **** p<0.0001.

### *Salmonella* MUC1-SiiE mediated invasion is dependent on α2,3-linked sialic acids

Next, we performed infection experiments to investigate the functional relationship between *Salmonella* SiiE, MUC1 and the different types of sialylated O-glycans. Bacterial invasion assays were performed with HT29-MTX WT and ΔMUC1 cultures and *Salmonella* WT and Δ*siiE* strains. Bacteria were pre-grown to EEP for optimal SiiE expression and incubated with the intestinal cultures for 1 hour, followed by killing of extracellular bacteria with gentamycin and the quantification of intracellular bacteria by colony counting. *Salmonella* WT invaded the HT29-MTX WT monolayers with high efficiency, amounting to 30% invasion of the initial inoculum. As previously reported, invasion was significantly reduced in the absence of MUC1 or SiiE (Fig. 6A). We determined motility of the two bacterial strains and found that the Δ*siiE* strain was as motile as the wild type in plate assays (Fig. S2). To address which sialic acid type was responsible for the SiiE-MUC1 interaction, the cellular monolayers were incubated with MALII to block α2,3-linked sialic acids or with SNA to block α2,6-linked sialic acids before addition of the bacteria. Addition of MALII significantly reduced invasion, to a similar level observed in ΔMUC1 monolayers while addition of SNA did not impact bacterial invasion (Fig. 6B). Addition of MALII or SNA to ΔMUC1 cells had no impact on invasion rates (Fig. 6B. Next, HT29-MTX cells were pretreated with the α2,3/6/8/9 or α2,3-specific neuraminidase for 3 hours prior to infection. Removal of all sialic acids or only the α2,3-linked sialic acids reduced the invasion to levels comparable to those in ΔMUC1 cells (Fig. 6C). Adding either type of neuraminidase to ΔMUC1 cells had no effect on invasion (Fig. 6C). To visualize the interaction of *Salmonella* with the α2,3-linked sialic acids, infection experiments were analyzed by confocal microscopy. In line with the results of the invasion assays, *Salmonella* expressing SiiE localized to MALII-positive areas and did not colocalize with SNA-positive areas (Fig. 6D-G). Together, our results demonstrate that α2,3-linked sialic acids that are part of α2,3-sTF and di-sTF structures on MUC1 are responsible for SiiE-mediated apical invasion of *Salmonella* into HT29-MTX cells (Fig. 6H).

**Figure 6.**
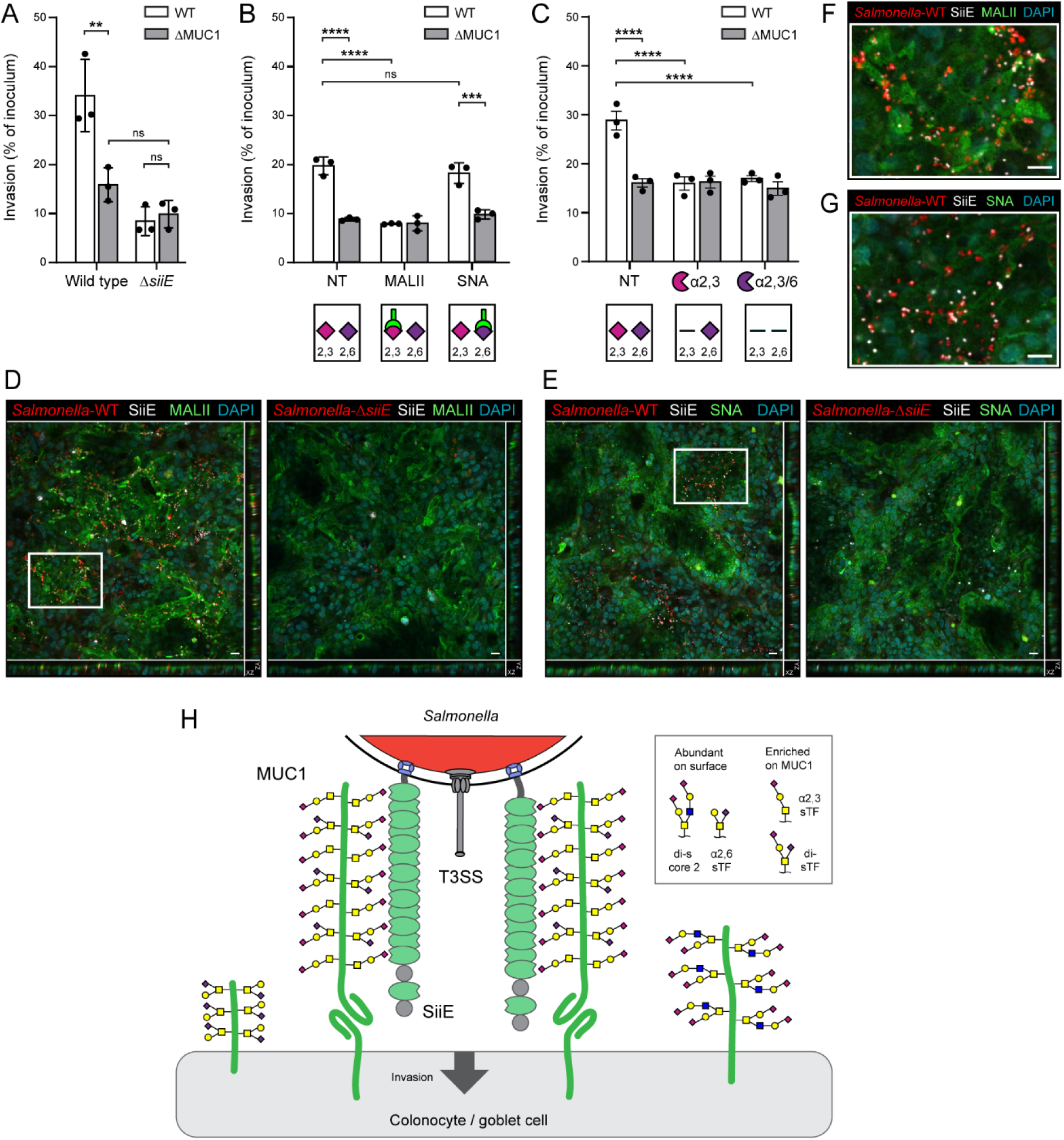
*Salmonella* invasion through the SiiE-MUC1 pathway is dependent on α2,3-linked sialic acids. A) Invasion assay of *Salmonella enterica* serovar Enteritidis WT and ΔSiiE in confluent HT29-MTX-WT and ΔMUC1 cultures. Bacteria were incubated with cultures for 1 hour and intracellular bacteria were quantified by gentamicin protection assay and colony counting. B) *Salmonella* invasion assay in combination with lectin blocking. Confluent HT29-MTX-WT and ΔMUC1 cultures were not treated (NT) or pre-incubated with MALII or SNA lectins for 30 minutes followed by incubation with *Salmonella*-WT bacteria for 1 hour. Intracellular bacteria were quantified by gentamicin protection assay and colony counting. C) *Salmonella* invasion assay in combination with neuraminidase treatment. Confluent HT29-MTX-WT and ΔMUC1 cells were not treated (NT) or treated with either α2,3-specific or α2,3/6/8/9 broad-specificity neuraminidases for 3 hours followed by incubation with *Salmonella*-WT bacteria for 1 hour. Intracellular bacteria were quantified by gentamicin protection assay and colony counting. D, E) Immunofluorescence of HT29-MTX-WT cultures incubated with mCherry-positive *Salmonella*-WT or *Salmonella*-Δ*siiE* (red) strained for SiiE (white) and DAPI (blue) in combination with MALII or SNA (green). F, G) Zoom in of *Salmonella-*WT images depicting combined *Salmonella*/SiiE/MALII and *Salmonella*/SiiE/SNA straining. H) Schematic model depicting *Salmonella* SiiE engagement of MUC1 through abundant α2,3-sTF and di-sTF glycan structures with α2,3-linked sialic acids. Graphs represent mean ± SD from three independent biological replicates. Statistical test: two-way ANOVA with Tukey’s correction. ns = not significant; ** *p*<0.01. *** *p*<0.005. **** *p*<0.001. Scale bars: 20 µm (D,E,F,G).

### MUC1 is expressed in the human colon and positive for α2,3-linked sialic acids

To further investigate the presence of MUC1 and different sialic acid linkages and their relevance for *Salmonella* invasion, we employed the InTESTine^TM^ model [44]. In this *ex vivo* model, human intestinal tissue is obtained from surgical resections, and freshly mounted in a custom-made transwell-like setup allowing apical infection and further analysis (Fig. 7A). Human ileum and colon tissue was obtained from the same donor and mounted in the InTESTine system within 2 hours after resection. Tissues were left uninfected or *Salmonella* was added to the apical compartment followed by incubation for 1 hour. Viability of the ileal and colonic segments was maintained for the duration of the experiment as measured by low levels of lactate dehydrogenase (LDH) in the supernatant (Fig. S3A). Uninfected and infected ileal and colonic segments were analyzed by immunofluorescence microscopy to investigate expression of MUC1 and the presence of α2,3-linked and α2,6-linked sialic acids (MALII/SNA). Our first observation was that MUC1 was detectable in the colonic segment, but not in the ileal tissue (Fig. 7B, C). This was in line with the SCS data that showed MUC1 expression in colonocytes but not enterocytes (Fig. 3A). We visualized that secreted mucus layer using Jacalin staining and observed a strong signal in the ileal tissues and some staining of the denser secreted mucus in the colonic tissue (Fig. S3B, C). In the ileal tissue, the goblet cells and glycocalyx stained positive for SNA but negative for MALII, suggesting that α2,6-linked sialic acids are highly abundant on secreted and transmembrane mucins in the ileum (Fig. 7B). By contrast, in the colonic tissue goblet cells and glycocalyx stained positive for MALII and negative for SNA, indicating the presence of α2,3-linked sialic acids (Fig. 7C). In the colon, MUC1 was most strongly detected in the intestinal crypts and co-localized with MALII staining (Fig. 7C, D).

**Figure 7.**
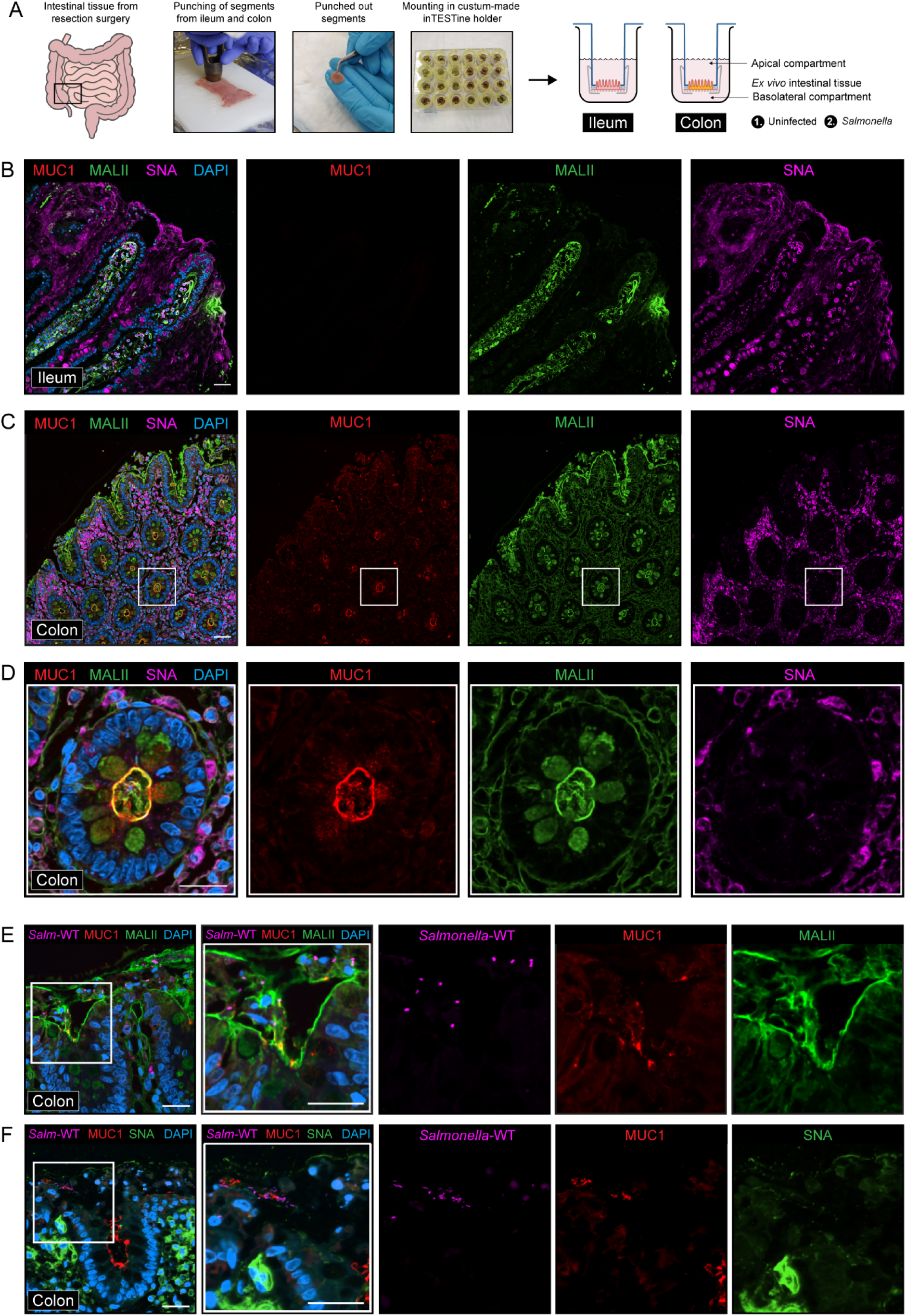
MUC1 expression and α2,3-sialylation is highly abundant in human *ex vivo* colon tissue. A) Workflow of the human InTESTine^TM^ *ex vivo* tissue model that allows analysis of and infection experiments with primary ileal and colonic tissues derived from resection surgery. Tissues are mounted in custom-made Transwell plates within 2 hours after resection and mucosal adaptation/recovery is allowed for 30 minutes. Cultures were fixed with 4% PFA, embedded in paraffin and sectioned for microscopy analysis. B) Uninfected human inTESTine distal ileal tissue stained for MUC1 (red), MALII (green), SNA (purple) and DAPI (blue). No MUC1 signal was detectable in ileal tissue slices. C) Uninfected human inTESTine proximal colonic tissue stained for MUC1 (red), MALII (green), SNA (purple) and DAPI (blue). MUC1 was highly expressed in colonic crypts and stained positive for MALII and negative for SNA. D) Zoom-in of colonic tissue from C. E, F) Colonic inTESTine tissues infected with *Salmonella enterica* serovar Enteritidis WT for 1 hour stained for MUC1 (red), MALII or SNA (green), *Salmonella* (FISH probe, magenta) and DAPI (blue). *Salmonella* can be detected in close proximity of MUC1 that is also stained by MALII. Scale bars: 50 µm (B,C,E,F) and 25 µm (zoom-in D-F).

Next, we investigated the localization of *Salmonella* in the infected *ex vivo* tissues using a previously reported specific FISH probe [47]. In ileal tissue, where no MUC1 was observed and the tissue stained highly positive for SNA, *Salmonella* WT could be detected close to the glycocalyx (Fig. S3D,E). In colonic tissue, *Salmonella* WT was detectable in close proximity to MUC1 that also stained positive for α2,3-linked sialic acids (MALII) (Fig. 7E). The proximity of *Salmonella* WT to MUC1 was also observed in regions where α2,6-linked sialic acids (SNA) were not detected (Fig. 7F). These *ex vivo* infection experiments corroborate the findings with the *in vitro* intestinal cultures regarding the importance of α2,3-sialylation of MUC1 for SiiE-mediated invasion. Furthermore, the InTESTine experiments demonstrate regional differences in MUC1 expression, sialic acid type and *Salmonella* invasion, and indicate the potential importance of the SiiE-MUC1 invasion pathway in the human colon but not in the ileum.

## Discussion

*Salmonella* bacteria are highly successful enteropathogens that have an array of virulence factors that allow survival, colonization, and invasion in the gastrointestinal tract. In this work, we investigated the molecular mechanisms of SiiE interaction with TM mucin MUC1 which mediates bacterial invasion at the apical intestinal epithelial surface. SiiE expression by *Salmonella* was highest in exponential growth phases, could be induced by oxygen shock, and levels of secreted SiiE are high in stationary cultures (Fig. 2). In different intestinal epithelial cell types, MUC1 is co-expressed with ST3GAL and ST6GALNAC sialyltransferases, suggesting the potential presence of both α2,3- and α2,6-linked sialic acids on MUC1 (Fig. 3). Colonic HT29-MTX cultures have both α2,3- and α2,6-linked sialylated core 1 and core2 O-glycan structures on their apical surface and immunoprecipitated MUC1 mostly carries the sialylated core 1 structures α2,3-sTF and di-sTF (Fig. 4, 5). *Salmonella* SiiE-mediated apical invasion was dependent on α2,3-linked sialic acids as demonstrated by specific removal or blocking of α2,3-linked sialic acids at the apical surface (Fig. 6). Finally, experiments with human *ex vivo* intestinal tissues revealed high expression of α2,3-sialylated glycan structures on MUC1 in the colon and not the ileum and localization of invasive *Salmonella* to these sites (Fig. 7).

Most *Salmonella* virulence genes are grouped in *Salmonella* Pathogenicity Islands (SPIs) SPI-1, SPI-2, SPI-3 and SPI-4 [48, 49]. SPI-1 is the most well-known and encodes a type III secretion system-1 (T3SS-1) and machinery essential for epithelial adhesion. *Salmonella* utilizes this T3SS-1 to inject effector proteins into host cells, triggering actin remodeling and bacterial uptake [30]. SiiE is encoded by the relatively understudied SPI-4 together with a type I secretion system (T1SS) involved in SiiE secretion [32]. SPI-4 is highly conserved between different Serovars including Enteritidis, Typhimurium and Typhi, underscoring its importance. However, the SPI-4 locus of *Salmonella* Typhi was reported to contain a stopcodon in SiiE leading to two predicted open reading frames [50, 51]. The available literature demonstrates that the contribution of SiiE to *Salmonella* pathogenesis depends on the host species. In orally infected 7-week-old pigs, a *S.* Enteritidis SPI-4 deletion mutant colonized the intestinal tract as efficiently as wild-type bacteria [52] and also in 2-week-old chicks, SPI-4 was not required for *S.* Typhimurium cecal colonization [53]. However, transposon insertion in SPI-4 compromises infection of calves [53] and SiiE contributes to colonization of the calf intestine by promoting invasion of enterocytes [54]. In mice, SPI-4 contributes to virulence of both *Salmonella* Enteritidis and Typhimurium following oral gavage [55], and the SPI-4 mutant of *S.* Typhimurium is fully avirulent during intraperitoneal infection of BALB/c mice [56]. In the mouse, it was demonstrated that also MUC13 plays a role during *S.* Typhimurium SiiE-mediated infection as SiiE interacted with MUC13 and MUC13 functions as a releasable decoy that limits bacterial epithelial invasion and barrier damage [57]. The current study together with publications from the group of Michael Hensel convincingly demonstrate that SiiE plays an important role in adhesion to and invasion of human polarized epithelial cells [33] and intestinal tissues (this study). It has been demonstrated the cooperation between the SiiE-encoding SPI-4 and SPI-1 that encodes the T3SS-1 is required for efficient penetration of human epithelial barriers [58]. Based on these findings in different host species, we propose that SiiE could be an important factor that contributes to host specificity and zoonotic potential of different *Salmonella* strains.

When enteropathogens arrive at the intestinal epithelium, oxygen levels increase compared to the lumen and this so-called “oxygen shock” is a known inducer of virulence genes [59]. We found that SiiE expression was indeed induced when bacteria transitioned from anaerobic growth to oxygen shock (Fig. 1). This increased expression presumably facilitates interaction with MUC1 on the apical surface. We also observed high SiiE expression during exponential growth, which is also in line with a previously published transcriptomic analysis of *Salmonella* in different infection-related growth conditions [34]. A striking observation was that a large portion of SiiE was secreted, especially at stationary phase. The function of secreted SiiE in the intestinal lumen is not completely clear, and one could hypothesize that it relates to interactions with mucins. However, it was demonstrated that secreted SiiE can have immune-suppressive functions. Secreted SiiE prevented antibody responses by selectively reducing the number of IgG-secreting plasma cells in the bone marrow of intraperitoneally infected mouse experiments with *Salmonella enterica* serovar Typhimurium [60]. This phenotype occurred through a laminin B1-binding section in SiiE and demonstrates that this giant adhesin has the capacity to interact with biological substrates other than O-glycans on mucin glycoproteins.

Unraveling the glycan specificity of bacteria-mucin interactions is an important aspect of mucosal infection biology. Using a MUC1 glycopeptide array, we found that secreted SiiE recognizes multiple classes of MUC1 O-glycans, including core 2, α2,3-sialylated, α2,6-sialylated, and disialylated glycoforms, with binding influenced by glycan position and density (Fig. 3). Although this microarray format reveals intrinsic SiiE glycan recognition preferences, it does not fully capture the higher-order avidity effects arising from interactions between the numerous lectin-like domains of SiiE and appropriately spaced O-glycans presented on native MUC1. For native glycosylated MUC1 on the intestinal epithelial surface, we demonstrated that α2,3-linked sialylation of O-glycans is required for SiiE-mediated invasion, whereas α2,6-linked sialylation, although recognized by SiiE, did not contribute to invasion (Fig. 6). Together, these findings demonstrate that glycan identity alone is insufficient to drive productive host recognition, rather, the spatial organization and multivalent presentation of glycan epitopes on MUC1 are critical determinants.

Through a combination of different approaches, we demonstrated the importance of the α2,3-linked sialic acid decoration of O-glycans on MUC1 for SiiE-mediated invasion, while α2,6-linked sialic acids were present but did not contribute (Fig. 6). For several of our assays, we made use of lectins that are widely used to study glycan linkages but also have some limitations. For example, MALII recognizes α2,3-sialylated termini but also 3-O-sulfated termini as both share a negatively charged glycan C3 substituent [61]. SNA is commonly described as α2,6-specific, but preferentially binds α2-6-sialic acid linked to Gal and has weaker binding to α2-6-sialic acid linked to GalNac (such as the sTn epitope) in glycan arrays [62–64]. Interesting tools in the glycan field include antibodies that are specific to glycan structures such as sTn [65], but their application is restricted to this single structure. As an alternative, glycan-recognizing affinity binders (GRABs), derived from catalytically inactive bacterial sialidases, can provide high-affinity recognition of sialic acids independent of linkage or, in some cases, with α2,3 specificity. While no α2,6-specific GRABs have been reported to date, this approach may offer a more precise strategy to detect defined sialic acid linkages [66].

Potential pathogenic specificity for α2,3-linked sialic acids on MUC1 seems to be a more common mechanisms as it was also observed for some *Helicobacter pylori* strains, which interact with sialyl Lewis x (sLe^x^) structures containing this linkage [19, 67]. In viral infections, sialic acid linkage is known to play a crucial role in determining host specificity [25, 68, 69]. Avian influenza predominantly binds to α2,3-linked sialic acids, which are the dominant sialic acid linked residues present in the respiratory and intestinal tracts of birds [70]. Human-adapted influenza strains preferentially bind to α2,6-linked sialic acids that are abundant in the human upper respiratory tract [71]. For bacteria, sialic acid linkage-specific interactions are less well studied.

The mucosal surface of the gastrointestinal tract is highly diverse, and its O-glycan composition is dependent on location, cell type, microbiome and inflammatory status [72]. Enhanced sialylation during inflammation can serve multiple purposes, for example protection of epithelial surfaces and dampening of immune responses [73, 74]. Our single-cell sequencing data showed that ST3GAL4 (α2,3) and ST6GALNAC1 (α2,6) are the primary transferases expressed in the intestinal tract (Fig. 2). In HT29-MTX cultures, ST3GAL1, ST3GAL4 and ST6GALNAC1 were the most highly expressed transferases (Fig 4A). ST3GAL1 has been described to be the main transferase responsible for sialylation of type 3 motifs in O glycans, while ST3Gal4 primarily targets LacNAc motifs [75]. Both α2,3-sialylated core-1 and core-2 structures were present in HT29-MTX cultures (Fig. 5), but the core-1 structures seemed to be enriched on MUC1 and potentially the target for *Salmonella* SiiE (Fig. 6). These structures align with the high expression of ST3GAL1 in the HT29-MTX cell line (Fig. 4A). However, activity of ST3GAL4 towards O-glycan core-1 structures has been suggested *in vitro* and under conditions of overexpression and therefore cannot be excluded [76]. During *H. pylori* infection, ST3Gal4 is responsible for generation of the sLe^x^ antigen that the bacteria can engage with and this enzyme was shown to be upregulated by the pro-inflammatory cytokine TNFα [77–79]. Also for *Pseudomonas aeruginosa* infection of lung epithelial cells, the upregulation of ST3Gal4 and subsequent increase in sLe^x^ structures was responsible for increased adhesion [79]. It is well known that *Salmonella* benefits from inducing an inflammatory state during infection [80, 81] and we can speculate that this inflammatory state results in an increase in ST3Gal1 and/or ST3Gal4 expression, with more α2,3-linked sialic acids and better SiiE adhesion to the epithelial surface and MUC1 in general.

ST6GALNAc1 is known to be responsible for aberrant truncated sTn glycan structures observed during inflammation and cancer [82–84]. However, in the intestine this enzyme also plays an important role in commensal homeostasis as ST6GALNAc1-mediated sialylation of MUC2 was shown to be protective and prevented degradation by mucinases [16]. In the single-cell sequencing data, ST6GALNAC1 expression was predominantly restricted to goblet cells, with little expression in colonocytes. In our HT29-MTX cultures, ST6GALNAc1 gene expression was high, but α2,6-sialylated O-glycans were not abundant, which perhaps relates to substrate competition with ST3GAL enzymes (Fig. 5). In the single-cell data, ST3GAL4 was highly expression in BEST4⁺ epithelial and Paneth cells (Fig. 2A). Consistent with previous studies, we observed that the ileum represents a predominantly α2,6-sialylated environment (Fig. 7) [85]. In the ileum, membranous epithelial (M) cells are a primary target for *Salmonella* invasion [86]. Our SCS data indicate substantial MUC1 expression in these cells (Fig. 3) and, in addition, M cells are known to display the α2,3-containing sialyl-Lewis A epitope [87]. It is therefore intriguing to speculate that M cells provide distinct α2,3-sialylated MUC1-positive surfaces and that SiiE could play an instrumental role in targeting bacteria to these sites to mediate invasion.

Using a MUC1 glycopeptide array, we found that SiiE could actually interact with different O-glycans and that binding to α2,3-sialylated O-glycans was relatively week. These data seem to be somewhat contradictory with the strong dependence on α2,3-sialylated O-glycans on MUC1 during invasion. However, in the full-length MUC1 that is expressed on the epithelial surface, the architecture of mucin VNTR regions and the spacing of O-glycan groups attached to serines and threonines within these regions can generate unique O-glycosylation patterns. Multiple interactions increase avidity and distinct patterns have been proposed to function as a specific “barcode” for receptor interactions [88]. In line with this concept, our research shows that the combination of the full length MUC1 backbone and its α2,3-sialylated O-glycans is essential for SiiE-mediated invasion. The presence of de-sialylated MUC1 after enzymatic removal or α2,3-sialylated O-glycans on other protein carriers do not confer invasion. We hypothesize that the spatial organization of α2,3-sialylated O-glycans along the 42 MUC1 VNTR domains as occurs in the colon most likely offers a highly favorable substrate for engagement by the 53 lectin-like domains of SiiE. It remains to be established if SiiE also has functional interactions with secreted MUC2 and/or TM mucin MUC13 in the human intestine. This question could be answered using novel glycan engineering techniques that make use of a HEK293 platform and tailored expression of mucin domains [12, 88–90]. Our study highlights the critical role of α2,3-linked sialic acids during *Salmonella* SiiE-mediated invasion through MUC1 and illustrates how *Salmonella* exploits host glycan structures that are present on specific cell types and locations for invasion.

## Acknowledgements

We thank Prof. Dr. Michael Hensel (Universität Osnabrück, Germany) for his generous gift of α-SiiE polyclonal rabbit serum. We thank Dr. John Hilkens for his support and providing α-MUC1 antibodies. We thank Filipa Marcelo for advice and support regarding the MUC1 glycan array. Microscopy was performed at the Center for Cell Imaging (CCI) of the Faculty of Veterinary Medicine, Utrecht University, and we thank Dr. Richard Wubbolts and Esther van ’t Veld for their expert advice. We thank Jordy van Angeren, from the Center for Proteomics and Metabolomics (CPM) at the Leiden University Medical Center (LUMC) for expert technical assistance.

## Data availability statement

All data discussed in this manuscript are available in the presented Figures and supplementary data.

### Author contributions

Conceptualization: KCAPG, JPMP, KS. Methodology: KCAPG, ALHE, HC, AD, BW, ASP, NH, JD, KS. Investigation: KCAPG, ALHE, HC, LZXH, AD, BW, JS, AK, ASP, JD. Visualization: KCAPG, KS. Supervision: KS, NH, JD. Writing original draft: KCAPG, KS. Review & editing: KCAPG, ALHE, HC, LZXH, AD, BW, JS, AK, ASP, JPMP, NH, JD, KS.

---

**Figure S1.**
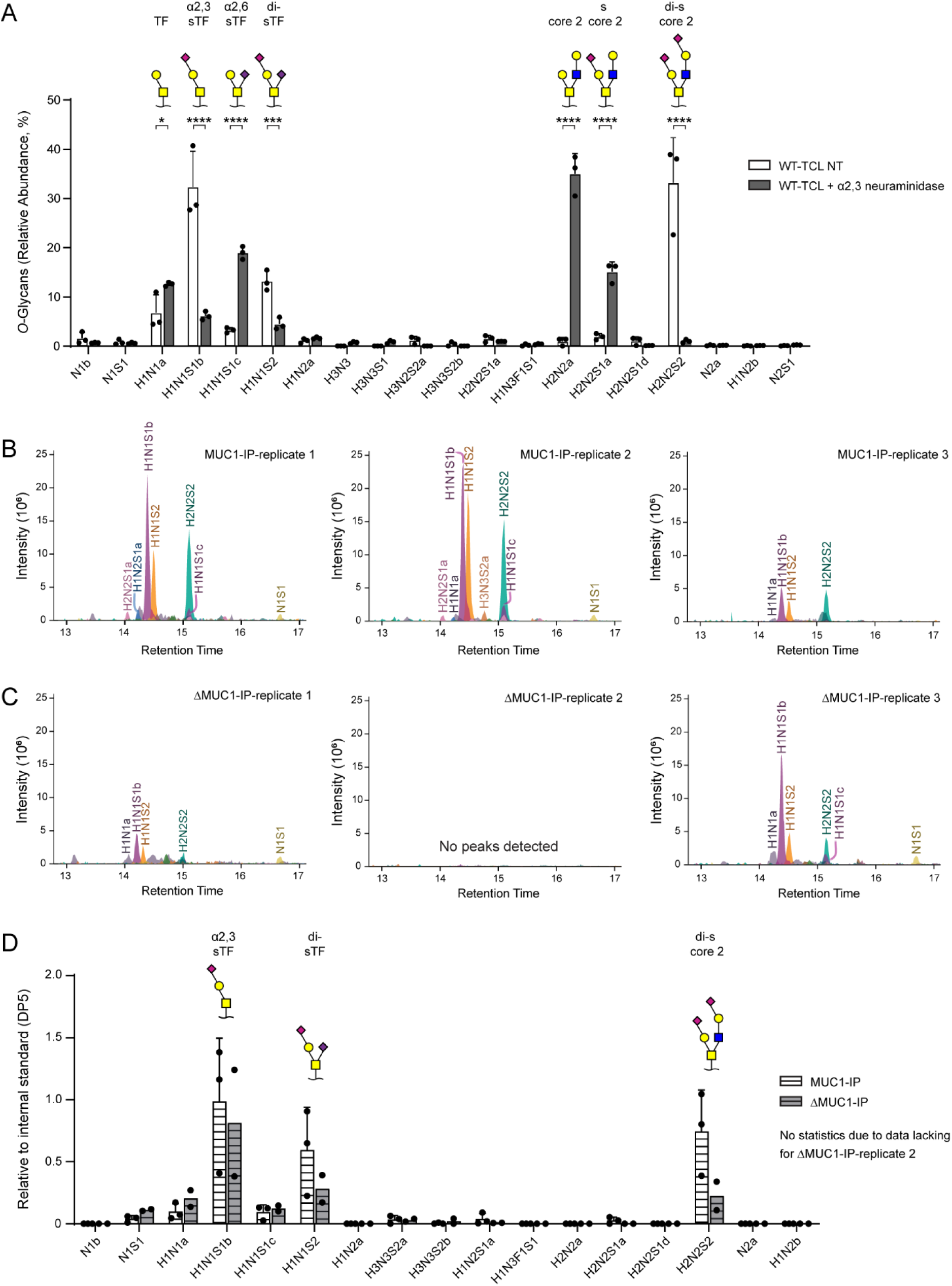
Additional *O*-glycomics analyses of HT29-MTX cultures and MUC1 immunoprecipitations. A) O-glycomics analysis of O-GalNAc structures present in HT29-MTX-WT total cell lysate (TCL) samples that were non treated (NT) or pretreated with α2,3-specific neuraminidase. The relative abundance of the identified O-GalNAc structures is plotted. The most abundant structures in the WT-TCL NT samples include α2,3-sTF (containing an α2,3-linked sialic acid), di-sTF (containing both an α2,3- and an α2,6-linked sialic acid), di-s core 2 (containing two α2,3-linked sialic acids) and in the WT-TCL + α2,3 neuraminidase samples TF, sTn sTn (containing a α2,6-linked sialic acid), core 2 and s core 2 are most abundant. H = hexose, N = *N*-acetylhexosamine, F = fucose, S = *N*-acetylneuraminic acid. a, b, c and d are added to the glycan name to distinguish LC-separated glycan isomers. B, C) Extracted ion chromatograms (EICs) of the most abundant O-GalNAc structures in MUC1-IP and ΔMUC1-IP samples. ΔMUC1-IP replicate 2 did not pass quality control resulting in no detection of peaks. D) Area under the curve data of MUC1-IP and ΔMUC1-IP data depicted in C, normalized against the DP5 internal standard. The most abundant O-GalNAc structures present in these samples included α2,3-sTF, di-sTF and di-score2. Graphs depict experimental results of two or three independent biological replicates and mean ± SD. Statistical test where applicable: two-way ANOVA with Tukey’s correction. * p<0.05; *** p<0.01. **** p<0.01.

**Figure S2.**
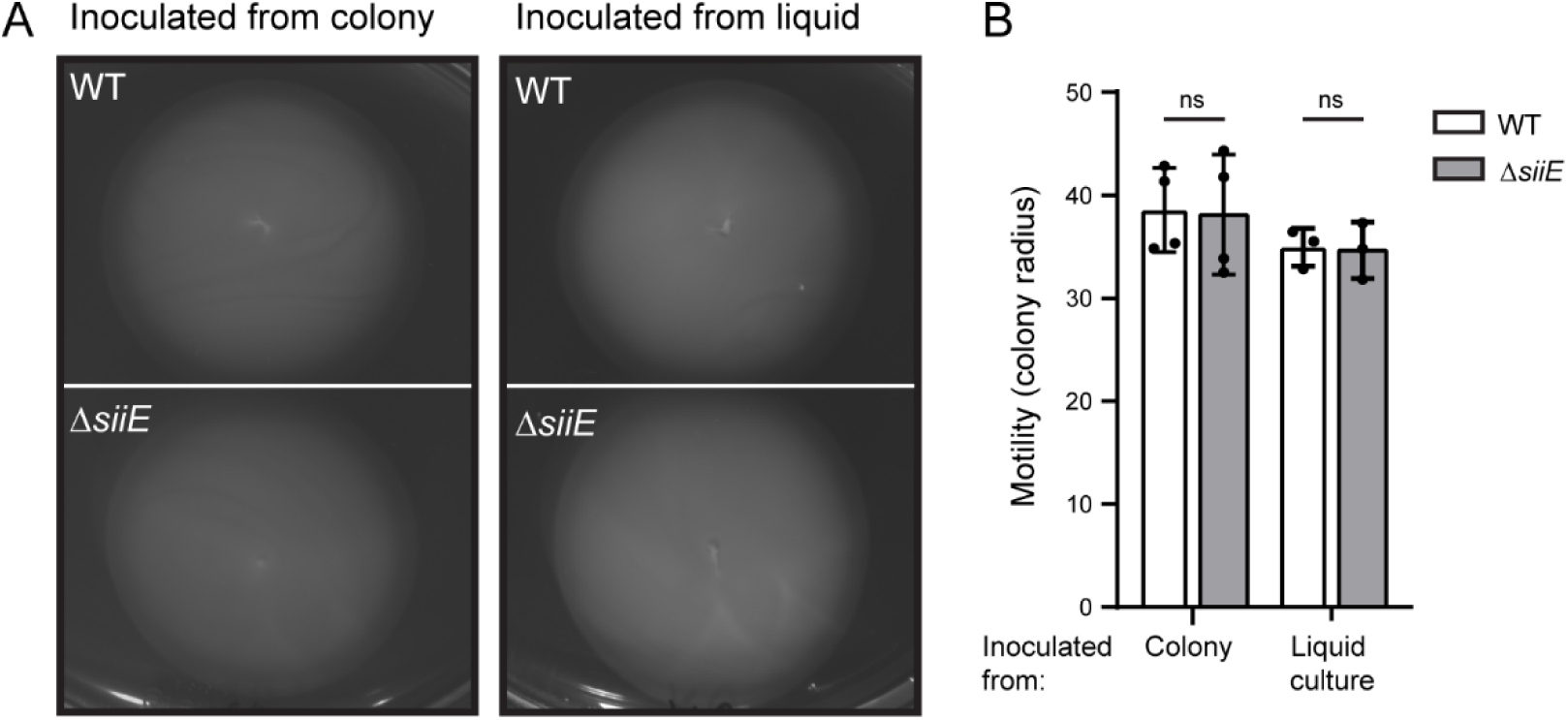
*Salmonella*-WT and *Salmonella-*Δ*siiE* bacteria have comparable motility in a plate assay. A) Representative images of motility agar plates of *Salmonella enterica* serovar Enteritidis WT and Δ*siiE* spotted from a colony a liquid pre-culture. B) Quantification of *Salmonella*-WT and *Salmonella*-Δ*siiE* motility. Graphs depict experimental results of three independent biological replicates with SD. Statistical test: two-tailed unpaired t-test. ns = not significant.

**Figure S3:**
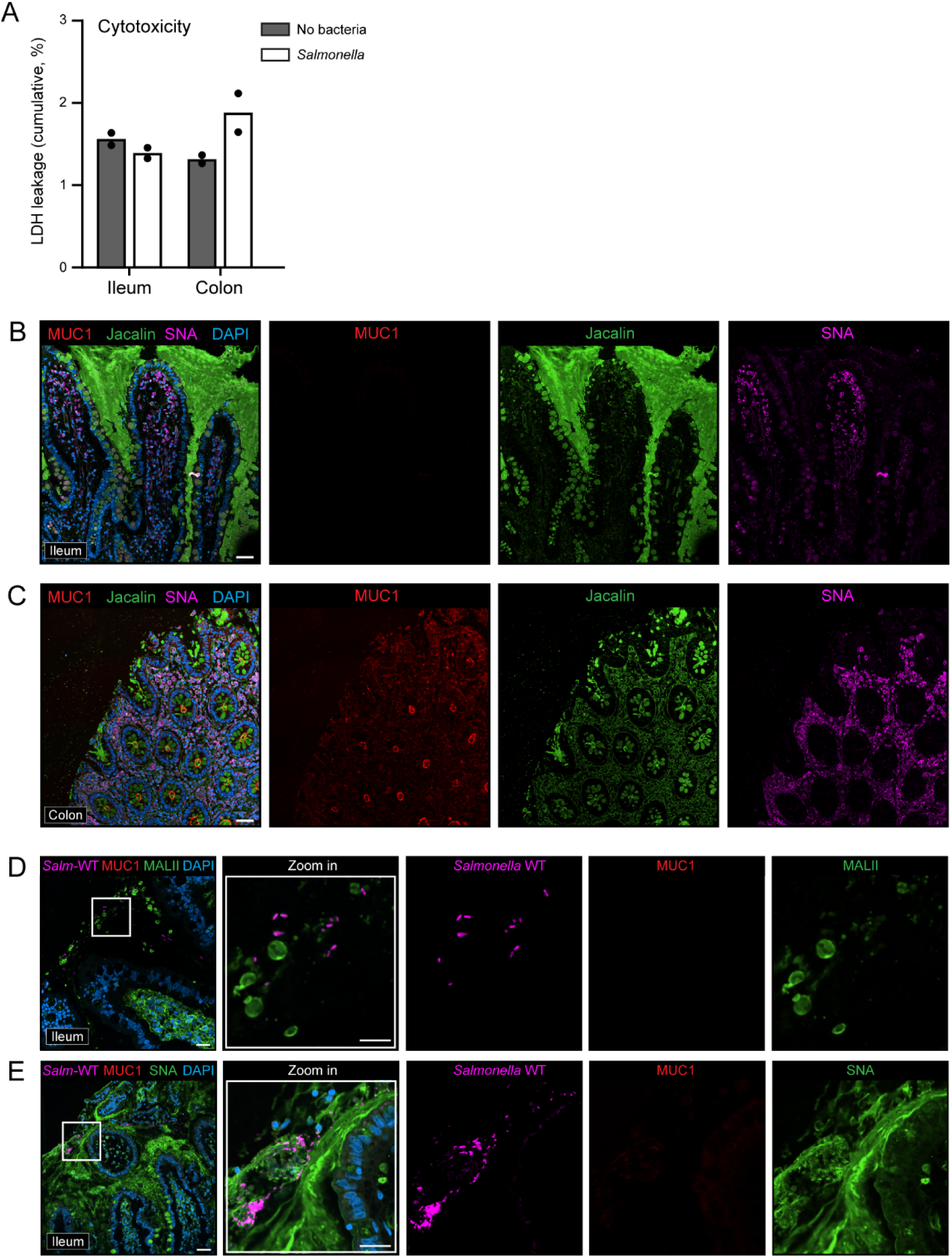
Additional data human inTESTine ex-vivo ileal and colonic tissues. A) LDH cytotoxicity measurements of human inTESTine™ *ex vivo* ileal and colonic tissues. Media from the apical and basal compartments were combined. Tissue viability was maintained throughout the experiment, with the LDH leakage below 10% per hour of the total LDH. Data represent measurements of two technical replicates from the same donor (*n* = 2 ileum, *n* = 2 colon). B) Uninfected human inTESTine distal ileal tissue stained for MUC1 (red), Jacalin (green), SNA (purple) and DAPI (blue). No MUC1 signal was detectable in the ileal tissues. C) Uninfected human InTESTine distal colonic tissue stained for MUC1 (red), Jacalin (green), MALII (purple) and DAPI (blue). D,E) Human InTESTine distal ileal tissue infected with *Salmonella enterica* serovar Enteritidis WT for 1 hour stained for MUC1 (red), MALII or SNA (green), *Salmonella* (FISH probe, purple) and DAPI (blue). Zoom-in on areas positive for *Salmonella*. Scale bars: 50 µm (B,C,D,E) and 25 µm (zoom-in D,E).

